# Collapse of late endosomal pH elicits a rapid Rab7 response via V-ATPase and RILP

**DOI:** 10.1101/2023.10.24.563658

**Authors:** R.J. Mulligan, M.M. Magaj, L. Digilio, S. Redemann, C.C. Yap, B. Winckler

## Abstract

Endosomal-lysosomal trafficking is accompanied by the acidification of endosomal compartments by the H^+^-V-ATPase to reach low lysosomal pH. Disruption of proper pH impairs lysosomal function and the balance of protein synthesis and degradation (proteostasis). We used the small dipeptide LLOMe, which is known to permeabilize lysosomal membranes, and find that LLOMe also impacts late endosomes (LEs) by neutralizing their pH without causing membrane permeabilization. We show that LLOMe leads to hyper-activation of Rab7 and disruption of tubulation and mannose-6-phosphate receptor (CI-M6PR) recycling on pH-neutralized LEs. Either pH neutralization (NH_4_Cl) or Rab7 hyper-active mutants alone can phenocopy the alterations in tubulation and CI-M6PR trafficking. Mechanistically, pH neutralization increases the assembly of the V_1_G_1_ subunit of the V-ATPase on endosomal membranes, which stabilizes GTP-bound Rab7 via RILP, a known interactor of Rab7 and V_1_G_1_. We propose a novel pathway by which V-ATPase and RILP modulate LE pH and Rab7 activation in concert. This pathway might broadly contribute to pH control during physiologic endosomal maturation or starvation and during pathologic pH neutralization, which occurs via lysosomotropic compounds or in disease states.

**Summary Statement:** Mulligan et al. show that V-ATPase assembly and RILP mediate hyperactivation of the small GTPase Rab7 when the late endosomal pH gradient collapses disrupting endosome tubulation and M6PR trafficking.

## Introduction

Lysosomes are the principal membrane-bound proteolytic compartment within cells and are the endpoints of degradative endocytic trafficking, phagocytosis, and autophagy. Progression of trafficking pathways to lysosomes is essential for maintaining proteostasis. Numerous pathologies are associated with impaired proteostasis, including disruption of lysosomal membranes leading to lysosomal membrane permeabilization (LMP). LMP is any perturbation that compromises integrity of the lysosomal limiting membrane, leading to leakage of luminal content into the cytosol (Wang et al., 2018), including protons (causing luminal pH neutralization), calcium and often cathepsins (Lakpa et al., 2021). LMP occurs in diseases including lysosomal storage disorders (Kirkegaard et al., 2010) and those with neurotoxic aggregates (Freeman et al., 2013; Dilsizoglu Senol et al., 2021; Flavin et al., 2017; Polanco and Götz, 2022). Still other genetic and neurodegenerative conditions impact maintenance of proper endosomal and lysosomal function, like pH, in the absence of LMP (Colacurcio and Nixon, 2016; Pescosolido et al., 2021). It is thus important to study how cells mechanistically respond to endosomal dysfunction, including repairing LMP or correcting pH disruption. Our understanding of cellular LMP responses has been greatly advanced by using the lysosomotropic dipeptide L-leucyl L-leucine-O-methyl-ester (LLOMe), which damages lysosomal membranes. Studies using LLOMe have uncovered rapid membrane repair mechanisms (within minutes), such as via ESCRT or PI4P/ATG2 (Bohannon and Hanson, 2020; Jia et al., 2020a; Niekamp et al., 2022; Radulovic et al., 2018; Radulovic et al., 2022; Skowyra et al., 2018; Tan and Finkel, 2022; Yim et al., 2022), as well as delayed lysosome-specific autophagy pathways (within hours) via galectins (Gal-3/8/9) and autophagy adaptors (Jia et al., 2020b; Papadopoulos et al., 2020; Zoncu and Perera, 2022; Yang and Tan, 2023). How pH disruptions play into such stress responses, however, is not fully elucidated.

Before reaching lysosomes, endocytosed cargo traffics through a series of endosomal compartments, including early (sorting) endosomes (EE), late endosomes (LE) and ultimately lysosomes (LYS). Maturation of LEs ensures endocytosed cargos encounter activated lysosomal hydrolases in increasingly acidified compartments before fusing with maximally degradative lysosomes. LE maturation is triggered by switching of Rab family proteins (i.e., Rab5 to Rab7). Active GTP-bound Rab7a (herein, Rab7) now engages Rab7 effector proteins (Huotari and Helenius, 2011), including the core retromer complex (Vps26/29/35) and RILP (Cantalupo et al., 2001; Progida et al., 2007). Rab7-Retromer has been implicated in retrieval of trafficking receptors like mannose-6-phosphate receptors (M6PRs) from LEs back to the trans-Golgi network (TGN), and for tubule generation necessary for recycling (Priya et al., 2015; Rojas et al., 2008; Seaman et al., 2009; Seaman et al., 2018). RILP links LEs to dynein motors for motility, but RILP/Rab7 complexes may also be involved in tubule formation from lysosomes (Mrakovic et al., 2012). These tubules likely require proper Rab7 GDP/GTP cycling for their formation (Markworth et al., 2021). Even though high-fidelity membrane trafficking through EEs/LEs is necessary for lysosomal function, studies regarding the impact of lysosomal disruption on endosomal trafficking or tubulation, and on regulation of Rab7 are lacking.

Maturation of LEs is further accompanied by decreasing intra-endosomal pH to that of the highly acidic lysosome (pH≈4.5). Cells establish and maintain endo/lysosomal pH through the endosome/lysosome-localized vacuolar H^+^-ATPase (V-ATPase), which is activated by the assembly of the cytosolic V_1_ complex with the membrane-embedded V_0_ domain (Collins and Forgac, 2020). Rab7 has been implicated in V-ATPase function and endosomal acidification by forming a ternary complex with its effector RILP and the V_1_G_1_ subunit of the V_1_-ATPase complex (De Luca et al., 2014) and by direct interaction with the V_0_a_3_ subunit in secretory lysosome trafficking in osteoclasts (Nakanishi-Matsui and Matsumoto, 2022). However, the precise connections between Rab7 activity, V_0_-V_1_ assembly and endosomal acidification are not well elucidated. Thus, it is unknown if luminal pH can coordinate cytoplasmic effector cascades like those of Rab7.

In this study, we sought to address the gaps in knowledge regarding how endosomal compartments critical for membrane trafficking (i.e., LEs) behave in the context of LMP. We report here that a pH-driven mechanism in otherwise undamaged LEs leads to V-ATPase- and RILP-mediated hyperactivation of the small GTPase Rab7, disrupting normal Rab7 tubulation and biosynthetic receptor trafficking. These findings suggest a Rab7-RILP-V-ATPase axis involved in pH modulation and pH-driven LE stress responses. We thus describe a novel LE stress response, likely relevant in LMP and pathologic pH neutralization.

## Results

### LLOMe disrupts lysosomes in NRK cells leading to expected LMP responses

In order to ask if and how endosomal organelles upstream of lysosomes were impacted in LMP, we treated normal rat kidney cells (NRK) and primary mouse embryonic fibroblasts (MEFs) with LLOMe. We first determined if these cells responded to LLOMe as previously described for other cell lines. We focused on key LMP responses including pH neutralization, direct membrane repair processes (i.e., ESCRT), and lysophagy (i.e., Gal3). We found each of these to occur in our cell lines, as pH-sensitive dextrans were de-quenched (pH neutralization, Fig. S1A-C, Video 1), ALIX was recruited following short LLOMe exposures (direct membrane repair, Fig. S1D-E, performed in MEFs due to antibody), and Gal-3 was recruited following long exposures (Fig. S1F-G). To further observe the kinetics of minimal LMP events, we loaded cells with Magic-Red Cathepsin B substrate (MR-B), a small peptide that fluoresces when cleaved by cathepsin B in degradative lysosomes. Unlike the larger dextrans, the fluorescent product of MR-B proteolysis is lost from degradative compartments in LLOMe indicating the presence of minimal LMP (Fig. S2A-D, NRKs).

Having established that LLOMe drives LMP as expected in NRK and MEF cells, we used a multi-marker immunostaining approach to ask whether LMP affected biosynthetic organelles [cis-Golgi and trans-Golgi Network (TGN), early (EE) and late (LE) endosomes] upstream of lysosomes (LYS). LLOMe treatment did not disrupt Golgi stacks, as markers for both the cis-Golgi (GM130) and trans-Golgi network (TGN38) were not altered (Fig. 1A-B). The appearance of EEs, marked by EEA1, was mildly, though not significantly (p= 0.0779), peripheralized in LLOMe (Fig. 1C, Fig. S2E-F). In contrast, we observed marked changes in Rab7 staining, a canonical LE protein, after 1 hour of LLOMe treatment (Fig. 1D), as compared to DMSO controls. The apparent volume of individual Rab7 compartments was increased whereas the total number of compartments was decreased in LLOMe (Fig. 1E-F). Staining was often punctate but seemed to decorate the edge of an enlarged compartment. Due to this discontinuous appearance, we wondered whether these were single compartments or clustering of smaller vesicles. Therefore, we performed ultrastructural analysis using transmission electron microscopy (TEM). TEM revealed the presence of numerous single-membraned vacuolar-like structures in LLOMe, rather than clusters of small vesicles (Fig. 1G), confirming our interpretation of singular enlarged Rab7-positive compartments. Further, measurements of the diameter of endosomes and lysosomes in electron micrographs revealed a shift toward larger structures in LLOMe (Fig. 1H, Fig. S3A). Whereas in DMSO conditions, the presence of enlarged endosomes was rare (Fig. S3B), in LLOMe conditions there were frequent examples of structures that are highly lucent (Fig. 1G, yellow, Fig. S3C,F) and those with dense, observable material within the lumen (Fig. 1G, black, Fig. S3D), suggesting that not all of the enlarged compartments are lysosomes. In keeping with this, we also observed double-membraned structures likely to be autophagosomes (Fig. S3E).

**Figure 1.**
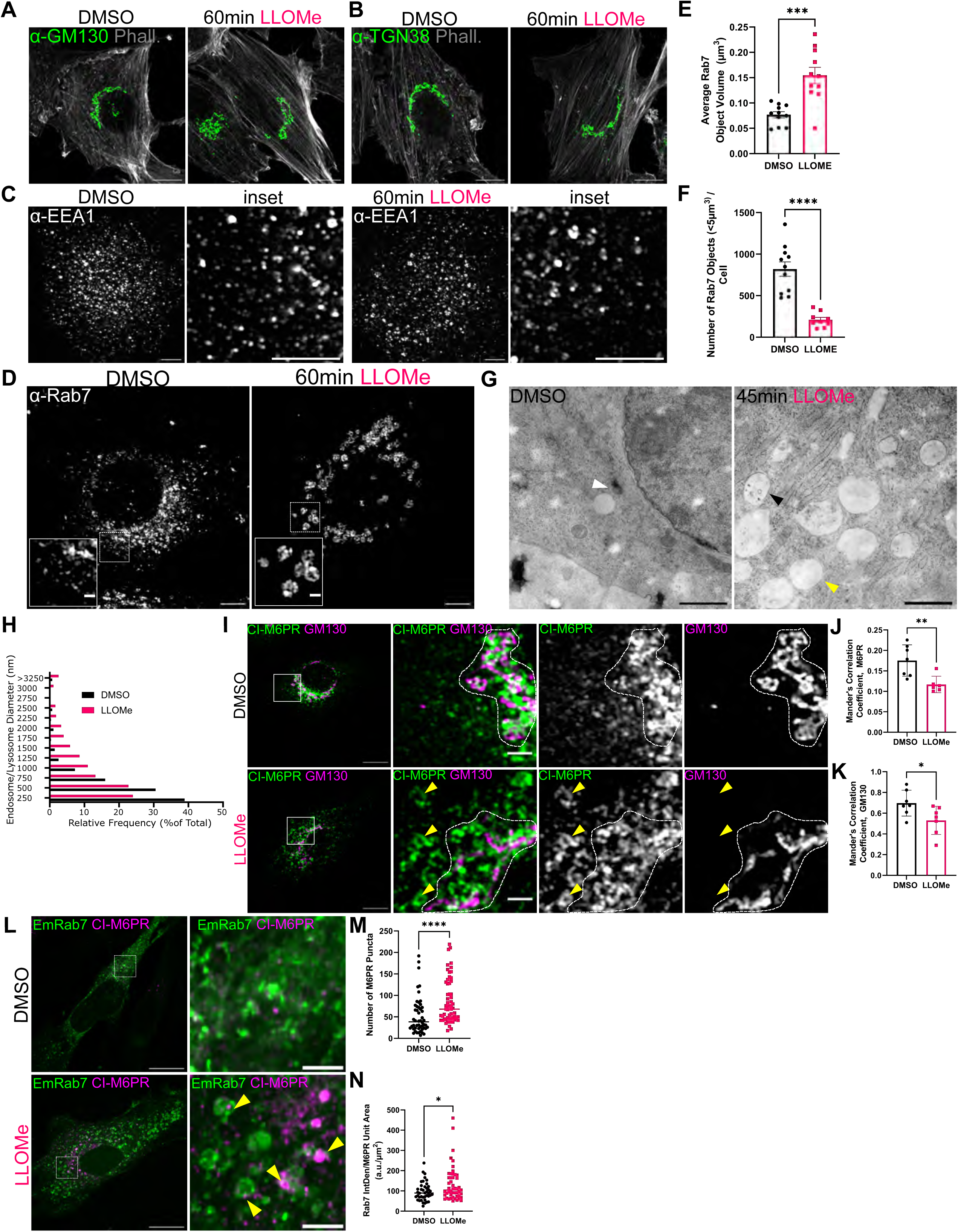
LLOMe treatments perturbs upstream Rab7-positive endosomes and M6PR-trafficking. **(A-C)** Normal rat kidney cells (NRK) were treated with DMSO or LLOMe (1mM, 1h)_and immunostained for (A) GM130 (cis-Golgi), (B) TGN38 (trans-Golgi network), and (C) early endosome antigen 1 (EEA1) (early endosomes) and imaged by Airyscan microscopy (scale= 5µm). Phalloidin staining is in white in A and B. **(D)** NRK were immunostained for Rab7 following DMSO or LLOMe (1mM, 1h) treatment (scale= 5µm, inset = 1µm) and imaged by Airyscan microscopy. Average Rab7 object volume **(E)** and number **(F)** per cell were determined and compared using Mann-Whitney-U (n= 55 cells, 3 independent experiments, ***p<0.001, ****p<0.0001, error bars= mean±SEM) **(G)** Representative transmission electron micrographs (TEM) of DMSO- and LLOMe-treated NRK cells (70nm sections, scale=2µm, n= 2 experiments). Highly electron-dense vesicle (lysosome), indicated with white arrowhead in DMSO (*left*). Single membraned, enlarged, lucent structures with (black) and without (yellow) luminal contents indicated in LLOMe (*right*). Additional examples in Fig. S3. **(H)** Relative frequency distribution of TEM diameters in DMSO and LLOMe, binned by 250nm (n=275(DMSO)/546(LLOMe) endosomes/lysosomes from 2 independent experiments) **(I)** NRK cells were immunostained for endogenous CI-M6PR (green) and GM130 (magenta) following DMSO or LLOMe (1mM, 1h) treatment (scale= 5µm, inset = 1µm), and imaged with Airyscan microscopy. GM130-positive Golgi area (dotted outline), peripherally mis-localized CI-M6PR (yellow arrowheads). **(J)** Mander’s correlation coefficients (MCC) for CI-M6PR intensity on GM130 space, and **(K)** GM130 intensity on CI-M6PR space compared using student’s t-test (n= 10 FOV, 3 independent experiments, *p<0.05, **p<0.01, error bars= mean±SD). **(L)** Confocal micrographs of primary MEFs transfected with Em-Rab7^WT^ and treated with either DMSO or LLOMe (1mM, 1hr) were fixed and stained for endogenous CI-M6PR (scale= 20µm, inset = 3µm). **(M)** Quantification of the total number of CI-M6PR puncta in (L). **(N)** Quantification of the intensity of Em-Rab7^WT^ signal on masked CI-M6PR area in (L). For M-N, conditions were compared using Mann-Whitney-U (n= 40 transfected cells per condition, 3 independent experiments) (*p<0.05, ****p<0.0001, lines = medians).

Since Rab7 compartments were enlarged in LLOMe, we wondered if the localization of cation-independent mannose-6-phosophate receptor (CI-M6PR), which traffics lysosomal cathepsins from the Golgi to endosomes but then recycles back from LEs to the TGN, was also disturbed. In control cells, we observed strong perinuclear colocalization between CI-M6PR and GM130, as expected. In contrast, CI-M6PR mis-localized in enlarged, more peripheral compartments in LLOMe (Fig. 1I) and co-localized less with GM130 (Fig. 1J,K). To test if these compartments were Rab7-positive, we expressed Emerald-Rab7^WT^ (Em-Rab7^WT^) in primary MEFs and immunostained for endogenous CI-M6PR. The peripheralized compartments containing mis-localized CI-M6PR were largely Rab7-positive (Fig. 1L; yellow arrowheads). Furthermore, peripheral CI-M6PR compartments were more abundant (Fig. 1M) and had more Rab7 on them, as compared to controls (Fig. 1N). Together, these results show that LLOMe-induced perturbation leads to rapid responses in Rab7+ compartments that retain CI-M6PR.

### Late endosomes rapidly respond to LLOMe

Given that Rab7 is present on multiple endosomal compartments, including transitioning EEs, LEs, and LYS, we investigated the identity of the LLOMe-responsive Rab7 compartments. We have previously distinguished EEs, LEs and LYS on the basis of multi-marker immunostaining and degradative capacity (Yap et al., 2018). We distinguish EEs from LEs based on the presence of EEA1, and LEs from LYS on the basis of degradative capacity, where LEs are degradatively (CatB or DQ-BSA) low and LYS are degradatively high (Fig. 2A). As compared to controls, transitioning EE-LEs (EEA1+Rab7+) in LLOMe-treated cells were not disrupted (Fig. S4A-C), despite the mild alterations in EE distribution (Fig. 1C, Fig. S2E-F). Interestingly, however, both LEs (Rab7+CatB*lo*: white arrowhead) and LYS (Rab7+CatB*hi:* yellow arrowhead) were enlarged following LLOMe treatment (Fig. 2B), consistent with our previous ultrastructural observations (Fig. 1G, Fig. S3). The co-localization of CatB with Rab7 was not changed (Fig. 2C) whereas LAMP1 staining was mildly enriched on Rab7 compartments following LLOMe treatment (Fig. 2D). LLOMe thus leads to phenotypic alterations in both LEs and LYS but not in transitioning EE-LEs. In order to determine the kinetics of Rab7+ responses to LLOMe, we carried out live imaging of NRK cells stably expressing Em-Rab7^WT^. We used DQ-BSA to distinguish degradative Rab7+ LYS from lowly degradative Rab7+ LEs. The apparent area of both LEs (Rab7+DQ*lo)* and LYS (Rab7+DQ*hi)* rapidly increased following LLOMe exposure (Fig. 2E-I, Fig. S4D, Video 2). These compartments continued to increase in size over the course of thirty minutes, but a significant size increase was apparent in less than ten (Fig. 2G,I). Interestingly, the Em-Rab7 signal expanded on both LEs and LYS with LLOMe, but the DQ-BSA within degradative LYS appeared contained within a subdomain of the enlarging LYS and did not fill the whole lumen. Similarly, endogenous CatB staining also often occupied only a subdomain within enlarged Rab7+ lysosomes (Fig. 2B). These data show that Rab7+ LEs are novel upstream responders to LLOMe.

**Figure 2.**
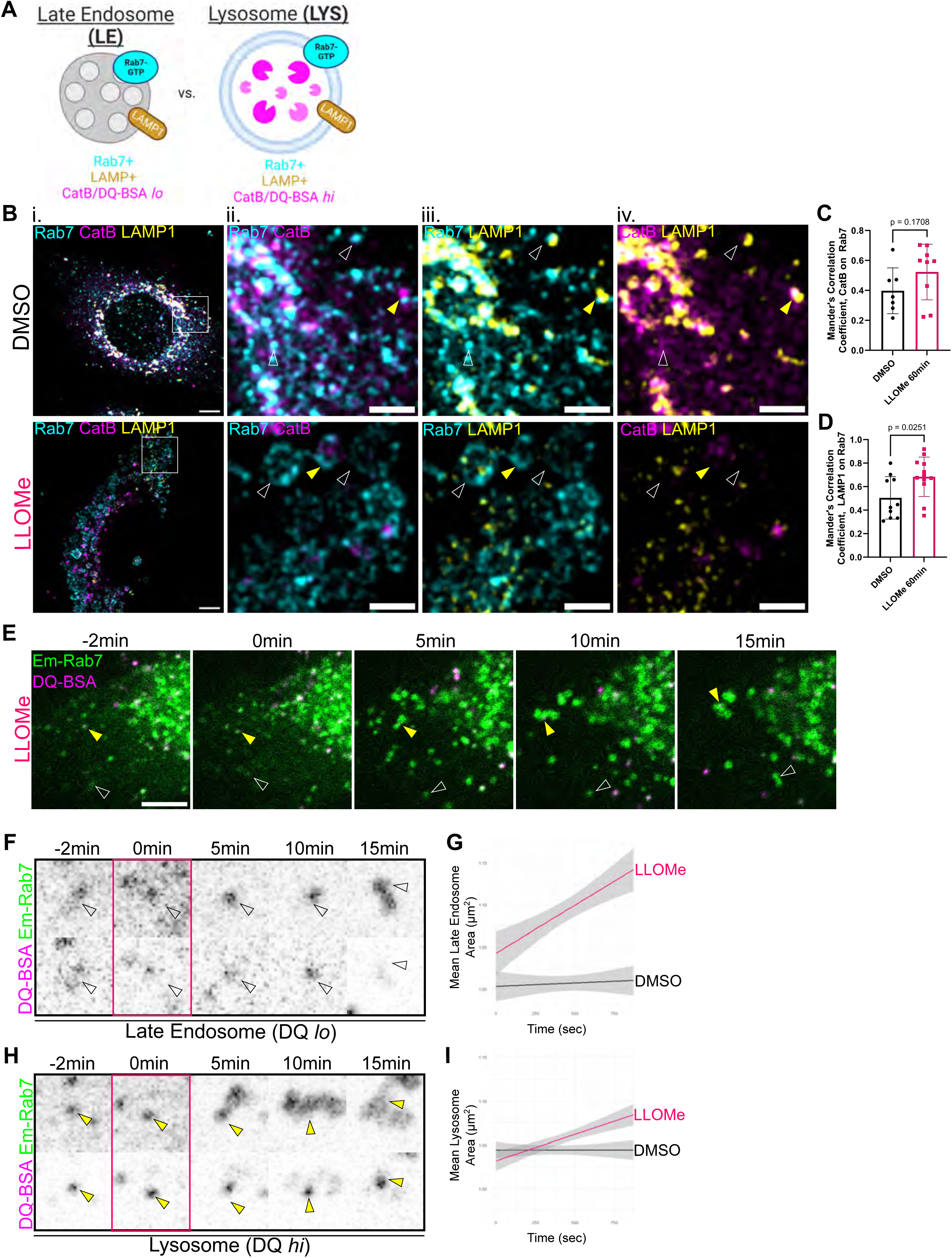
Rab7-positive late endosomes rapidly respond to LLOMe. **(A)** Definitions of late endosomes (LE) versus lysosomes (LYS). Both are positive for Rab7 (cyan) and lysosomal associated membrane protein 1 (LAMP1) (yellow) but differ on degradative ability (CatB*lo*/DQ-BSA*lo* versus CatB*hi/*DQ-BSA*hi*, magenta). **(B)** NRK cells were immunostained for Rab7, LAMP1, and lysosomal hydrolase cathepsin B (CatB) following DMSO or LLOMe (1mM, 1h) treatment (scale= 5µm, inset = 2µm) and imaged by Airyscan microscopy. Representative LE (white arrowhead) and LYS (yellow arrowhead) are indicated. MCC for CatB on Rab7 space (p = 0.1708) **(C),** and LAMP1 on Rab7 space (p = 0.0251) **(D)** were determined and compared using student’s t-test (n= 10 FOV per condition, 3 independent experiments, error bars= mean±SD). **(E)** EmRab7^WT^-transfected NRK cells loaded with 5µg/mL DQ-BSA overnight (Video 2), were imaged live in the presence of 1 mM LLOMe for 15mins. Still frames are shown. LEs (Rab7+DQ*lo*, white) and LYS (Rab7+DQ*hi*, yellow) are indicated. Single channel insets of a representative LE **(F)** and LYS **(H)** are shown. Mean area of DQ*lo* LEs **(G)** and DQ*hi* LYS **(I)** per cell were quantified over time (n= 10 (DMSO) and 9 (LLOMe) movies).

### Rab7 is hyper-activated following LLOMe treatment

We next sought to elucidate what could lead to the changes observed in Rab7+ compartments. LLOMe-mediated Rab7+ endosome enlargement (Fig. 1) is reminiscent of the accumulation of constitutively active (CA) Rab7^Q67L^ on enlarged endosomes (Bucci et al., 2000). CA-Rab7 is GTP-bound, which binds strongly to effectors. Thus, we sought to test whether LLOMe alters Rab7 nucleotide binding via interaction with an immobilized effector. To do so, we expressed GFP-tagged Rab7^WT^ or GFP-CA-Rab7^Q67L^ in COS-7 cells, treated with DMSO or LLOMe, and immunoprecipitated lysates with a GST-fusion protein containing the Rab7-binding domain of RILP, to specifically capture active Rab7-GTP. GFP-CA-Rab7^Q67L^ was used as a positive control to provide the maximum possible effector binding. Lysates from LLOMe-treated cells revealed a moderate but consistent increase (20%) in bound GFP-Rab7, as compared to DMSO-treated controls (Fig. 3A-B).

**Figure 3.**
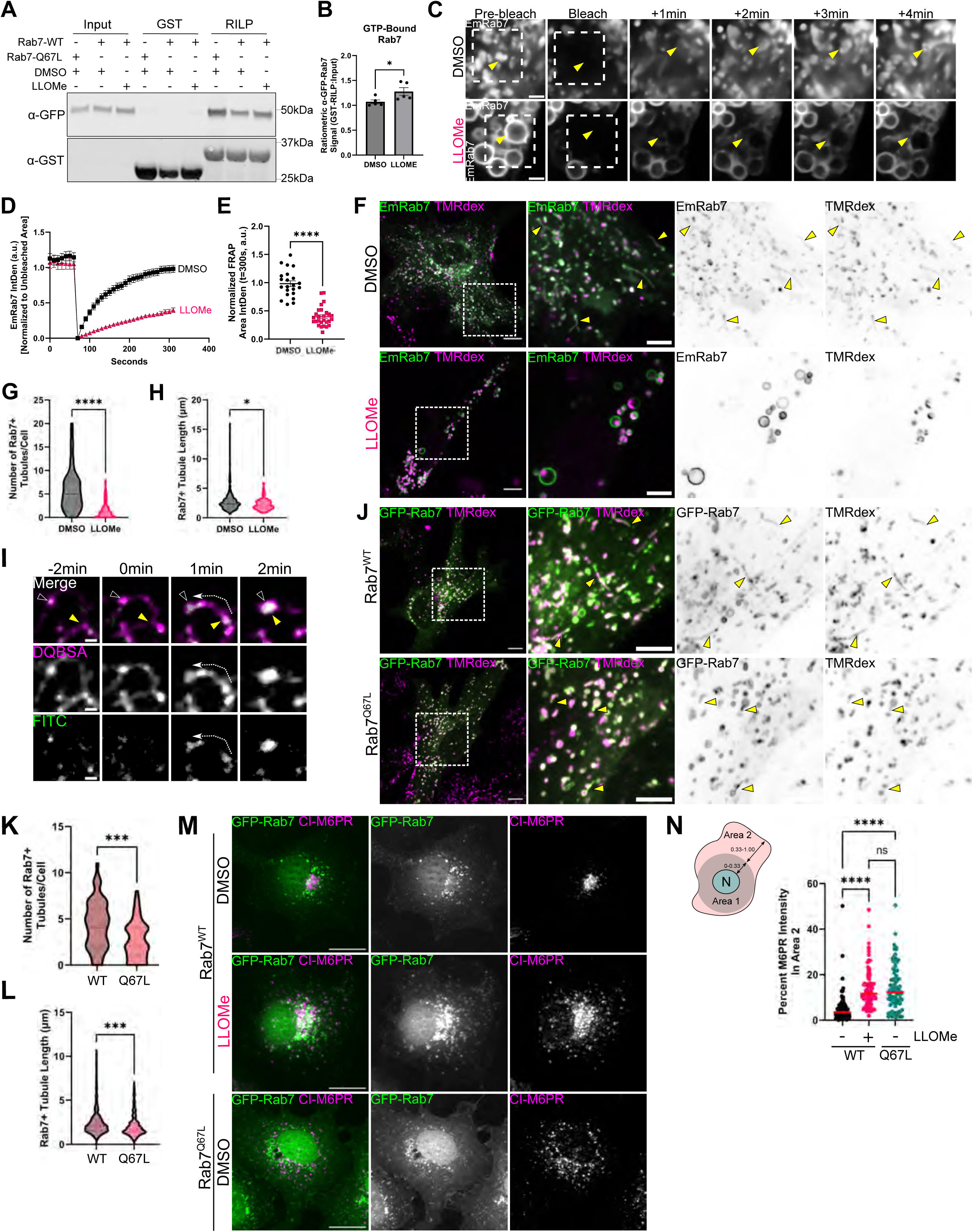
LLOMe-driven hyper-activation of Rab7 is sufficient to disrupt endosomal tubulation and CI-M6PR trafficking. **(A)** Representative immunoblot of GST or GST-RILP immunocaptured lysates from GFP-Rab7 transfected COS-7 cells treated with either DMSO or LLOMe (1mM, 1hr). GFP-Rab7^Q67L^ was used as a positive control for maximum GTP binding. **(B)** Quantified RILP-bound GFP-Rab7 in DMSO- and LLOMe-conditions by densitometry using student’s t-test (n= 5 independent experiments, p = 0.0433, error bar = mean±SEM). **(C)** Representative stills from confocal fluorescence recovery after photobleaching (FRAP) live imaging (Video 3) of Em-Rab7^WT^ expressing cells under DMSO (10mins, n=23 cells) and 1mM LLOMe (10mins, n=27 cells) conditions. Bleach area indicated by white box (33.0625µm^2^) with representative Rab7+ endosomes indicated by yellow arrowhead (scale = 2µm). **(D)** Normalized FRAP recovery curves were plotted over time. **(E)** Mean FRAP area intensity at t=300sec (Y_max_) was compared using student’s t-test (n= as in (C) from 3 independent experiments). **(F)** Confocal micrographs of Em-Rab7^WT^ transfected primary MEFs loaded with 100µg/mL tetramethyl-rhodamine dextran (TMRdex; magenta) for 2hr, prior to treatment with DMSO or 1 mM LLOMe for 45min. Em-Rab7^WT^ tubules (green) = yellow arrowheads (scale= 10µm, inset= 5µm). **(G)** Quantification of the number of Em-Rab7^WT^+TMRdex+ tubules in (F), compared using Mann-Whitney-U (n=57 (DMSO) and 50 (LLOMe) cells, 3 independent experiments). **(H)** Quantification of the length of Em-Rab7^WT^+TMRdex+ tubules from conditions in (F), compared using Mann-Whitney-U test (n=331 (DMSO) and 64 (LLOMe) tubules, 3 independent experiments). **(I)** Representative stills from live imaging (Video 4) of LLOMe-treated MEFs loaded with 5µg/mL DQ-BSA overnight and 100µg/mL FITC-dextran for 30min prior to imaging. Yellow arrowhead=terminal end of a tubule collapsing into its source endosome (white arrowhead). **(J)** Confocal micrographs of GFP- Rab7^WT/Q67L^ transfected primary MEFs loaded with 100µg/mL TMRdex for 2hr (scale= 10µm, inset= 5µm). **(K)** Quantification of the number of GFP-Rab7^WT/Q67L^TMRdex+ tubules compared using student’s t-test (n=62 (WT), 59 (Q67L) cells, 2 independent experiments). **(L)** Length quantification of GFP-Rab7^WT/Q67L^TMRDex+ tubules, compared using Mann-Whitney-U (n=266 (WT) and 159 (Q67L) tubules, 2 independent experiments). **(M)** Widefield micrographs of GFP-Rab7^WT^ (DMSO or LLOMe 1 mM, 1 h) and GFP-Rab7^Q67L^ (DMSO) transfected COS-7 cells, immunostained for endogenous CI-M6PR (scale = 20µm). **(N)** Binned CI-M6PR objects and measured for intensity of CI-M6PR in Area 2 compared using Kruskal-Wallis test (n=83 (WT-DMSO), 88 (WT-LLOMe), and 78 (Q67L-DMSO) cells, 3 independent experiments) (n.s. = non-significant, *p<0.05, ***p<0.001, ****p<0.0001).

To further validate that Rab7 was hyperactivated following LLOMe, we performed fluorescence recovery after photobleaching (FRAP) experiments to observe Rab7 membrane dynamics. Active Rab7-GTP is strongly associated with endosomal membranes compared to GDP-Rab7 and previous work reported slowed exchange of CA-Rab7^Q67L^ on endosomes via FRAP (McCray et al., 2010). We carried out FRAP on Rab7+ compartments in NRK cells stably expressing Em-Rab7^WT^ following treatment with either DMSO or LLOMe for 10 minutes (Fig. 3C, Video 3). Control FRAP experiments led to 50% recovery of Rab7 fluorescence within one minute and complete recovery to pre-bleach intensity within four minutes (Fig. 3C-D, black line). LLOMe treatment led to slower (Fig. 3C-D, pink line) and incomplete fluorescence recovery, which was significantly different than that of controls (Fig. 3E, pink). Together, these data suggest that LLOMe leads to increased GTP-bound and hyper-stabilized Rab7 on enlarged endosome/lysosomes.

A recent report demonstrated that Rab7 needs to be inactivated for some tubulation events to take place (Bhattacharya et al., 2023). Since we observe hyperactivated Rab7 after LLOMe treatment, we investigated whether tubulation of Rab7+ compartments was impacted. We loaded primary MEFs transfected with Em-Rab7^WT^ continuously with tetramethyl-rhodamine dextran (TMRdex) for 2 hours to populate all endosomal compartments and then treated with either DMSO or 1mM LLOMe for 45 minutes. Live imaging of Em-Rab7^WT^+/TMRdex+ compartments in DMSO revealed many tubules emanating from Rab7+ compartments (Fig. 3F). Interestingly, the exposure to LLOMe led to a significant reduction in the number of Em-Rab7^WT^+/TMRdex+ tubules per cell (Fig. 3F-G). The small number of tubules still forming in LLOMe-treated conditions were shorter and had a reduced range of sizes (Fig. 3H). To characterize acute tubule dynamics in relation to pH neutralization following LLOMe, we performed live imaging of cells loaded with DQ-BSA and FITC-dextran. DQ-BSA containing tubules collapsed within 2 minutes of LLOMe exposure, which was temporally linked to de-quenching of FITC dextran (i.e., increased luminal pH) (Fig. 3I, Video 4). These data suggest that Rab7+ tubules that contain luminal cargoes are reduced in the presence of LLOMe. Thus, these observations might explain both the CI-M6PR mis-localization and the increased Rab7 compartment size.

We next asked whether Rab7 hyper-activation alone was sufficient to lead to the observed deficits in CI-M6PR trafficking and Rab7+ endosome tubulation. To do so, we overexpressed CA-Rab7^Q67L^ and assessed endosomal tubulation in primary MEFs, and endogenous CI-M6PR distributions in COS-7 cells, respectively. Rab7^Q67L^ overexpression alone was sufficient to significantly reduce the number and length of Rab7+/TMRdex+ tubules as compared to Rab7^WT^ (Fig. 3J-L). Similarly, Rab7^Q67L^ overexpression led to CI-M6PR mis-localization akin to LLOMe treated Rab7^WT^ cells, as CI-M6PR staining was significantly increased in the peripheral two thirds of cell areas, as compared to control (Fig. 3M-N). Together, these data suggest that hyper-activation and hyper-stability of Rab7 on endosomes following LLOMe is sufficient to drive endosomal tubulation deficits and receptor mis-trafficking.

### LLOMe does not rupture Rab7-positive late endosomes but does neutralize their pH

We next sought to uncover the signal following LLOMe-induced perturbation that could drive changes in Rab7. Given the lysosomotropic weak base properties of LLOMe, more LLOMe will concentrate in more acidic, mature compartments compared to earlier more neutral compartments (Thiele and Lipsky, 1990; Uchimoto et al., 1999). Since LEs are relatively acidic (∼pH=5.5), Rab7+ LEs could concentrate enough LLOMe to damage membranes, explaining common phenotypes of LE and LYS. Alternatively, it is possible that LLOMe only damages membranes of highly acidified LYS but not of less acidified LEs. In this case, two alternative hypotheses are that Rab7+ LE changes result from signaling cascades via leaking LYS content (e.g., Ca^2+^) or that Rab7+ LEs are undergoing a direct LLOMe-mediated stress response in the absence of LMP.

To evaluate these possibilities, we first determined if LLOMe causes membrane permeabilization of LEs. We stained for endogenous ALIX (as in Fig. S1) combined with Rab7 and partitioned LEs from LYS by presence of CatB. Following 5 minutes of 1mM LLOMe exposure, ALIX was markedly recruited to organelles as compared to controls (Fig. 4A-B; yellow). We are able to observe both damaged (ALIX+) and undamaged (ALIX-) LYS (Rab7+CatB*hi*) and LEs (Rab7+CatB*lo*) (Fig. 4C). Quantification of damaged species revealed >50% of LYS recruited ALIX within the first 5 minutes of LLOMe exposure (Fig. 4D) while only 15% of LEs did (Fig. 4E). Of the total LLOMe-induced ALIX+ compartments, 67.82% were LYS while only 32.18% are LEs (Fig. 4F). Thus, these data suggest that LYS are the predominant species with membrane damage following LLOMe, while LE membranes are largely preserved. To see if Rab7+ endosomes would progress to stages of extensive LMP involving lysophagy, we co-immunostained treated NRK cells for Rab7 and Gal3. We observed minimal overlap of Rab7 compartments with Gal3, indicating the majority of LEs do not develop extensive damage (Fig. 4G). These data suggest that phenotypic changes in Rab7 are not dependent on endosomal membranes being physically damaged.

**Figure 4.**
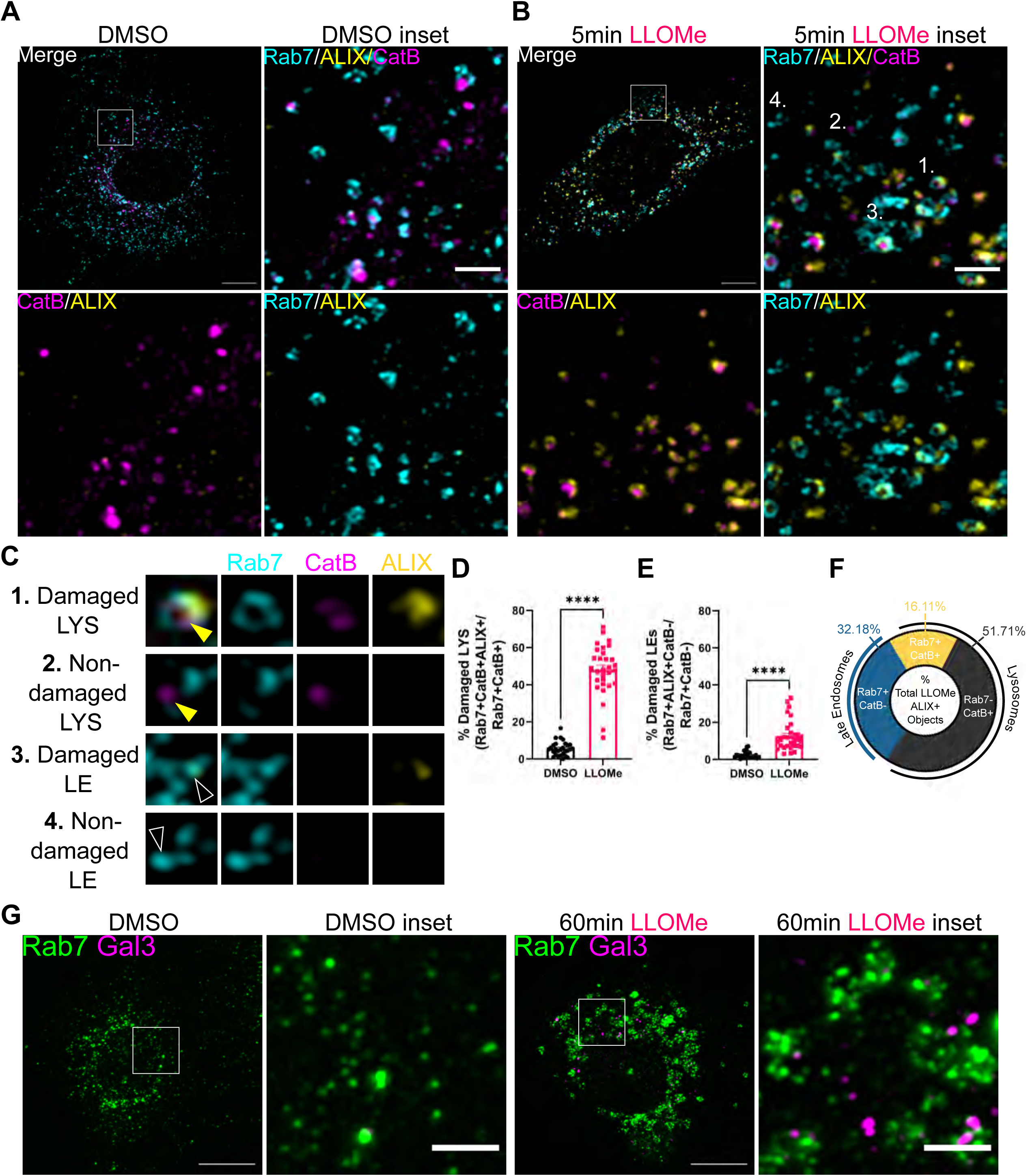
Most Rab7-positive late endosomes do not sustain membrane rupture after LLOMe. Airyscan micrographs of primary MEFs immunostained for endogenous Rab7 (cyan), ALIX (yellow), and CatB (magenta) following **(A)** DMSO or **(B)** 5-minute LLOMe treatment (scale= 10µm, inset = 2µm). Representative damaged LYS (1; CatB+ALIX+), non-damaged LYS (2; CatB+ALIX-), damaged LE (3; CatB-ALIX+), non-damaged LE (4; CatB-ALIX-) in **(C)**. The percentage of total ALIX+ LYS **(D)** and LE **(E)** objects were quantified and compared using Welch’s t-test (n= 30 FOV, 3 independent experiments, ****p<0.0001. Error bars=mean±SEM.). **(F)** The proportion of ALIX+ objects in LLOMe that are LE vs LYS. **(G)** Airyscan micrographs of NRK cells treated with DMSO or LLOMe (1mM, 1h) and immunostained for endogenous Rab7 and Gal3 (scale= 10µm, inset = 2µm). Rab7 intensity levels were artificially matched for viewing.

We next investigated the alternative hypothesis that luminal content from damaged LYS might cause phenotypic changes in size and number of Rab7 compartments in response to LMP. In particular, local Ca^2+^ release from LYS after LLOMe is required for both ESCRT-dependent and -independent repair mechanisms (Niekamp et al., 2022; Radulovic et al., 2022; Skowyra et al., 2018; Tan and Finkel, 2022; Yim et al., 2022). Cytosolic Ca^2+^ release can also elicit signaling through adaptor proteins (Clapham, 2007) that have profound impacts on membrane trafficking (Chadwick et al., 2021), and thus might drive downstream changes in Rab7+ LE size and number. However, local Ca^2+^ chelation with BAPTA-AM in LLOMe, or stimulation of Ca^2+^ release with TRPML1 agonism were not sufficient to drive Rab7 phenotypic changes (Fig. S5A-E, Video 5). Together, these data suggest that Ca^2+^ release, the most likely ionic signaling molecule for LYS to LE communication, is not responsible for observable changes in Rab7+ LEs.

Finally, we investigated the hypothesis that Rab7+ compartments were undergoing a direct LLOMe-mediated stress response independent of membrane damage. Given the weak base properties of LLOMe, we posited that LLOMe treatment might lead to direct pH neutralization of LEs without membrane damage and ALIX recruitment. To test this, we loaded cells with DQ-BSA overnight and then acutely loaded pH-sensitive Oregon Green-dextran (OGdex) for 1hr (to populate all endocytic compartments) before live imaging of cells exposed to DMSO or LLOMe (Fig. 5A). As expected, LLOMe treatment led to rapid de-quenching of OGdex in LYS (DQ*hi)*, indicative of pH neutralization (Fig. 5B-C,E, yellow arrowhead, Video 6). Similarly, OGdex was also de-quenched in other endocytic compartments (DQ*lo)* following LLOMe treatment (Fig. 5B,D,F, empty arrowhead, Video 6). Overall, these data suggest that while LLOMe does not extensively damage LEs as compared to LYS, it accumulates enough to neutralize their pH.

**Figure 5.**
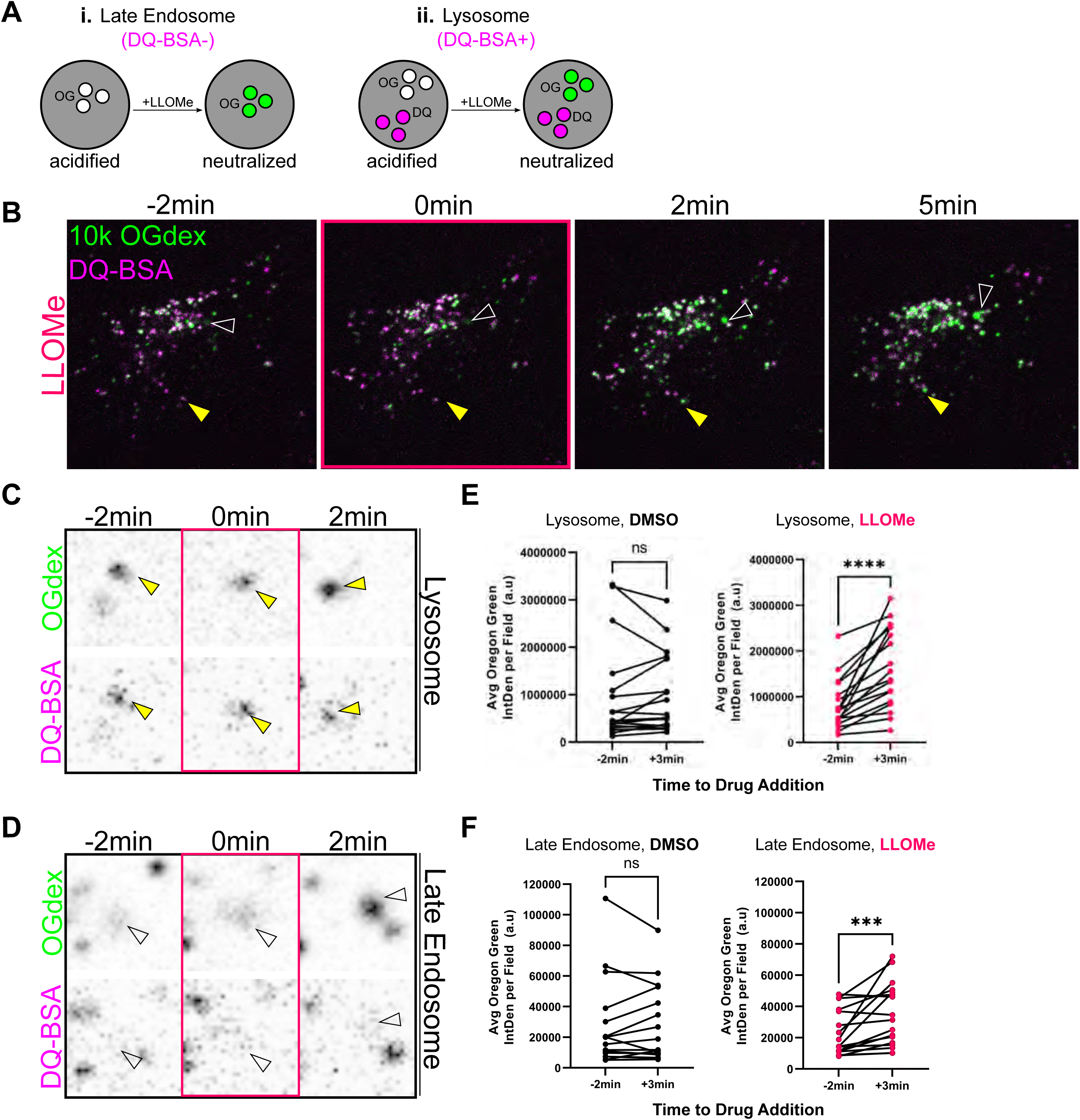
LLOMe leads to pH collapse of Rab7-positive late endosomes and lysosomes. **(A)** pH disruption de-quenches OGdex fluorescence in DQ*lo* LEs and DQ*hi* LYS. **(B)** Frames from live spinning disk microscopy (Video 6) of NRK cells preloaded with 5µg/mL DQ-BSA and then loaded with 100µg/mL OGdex for 1hr, followed by live LLOMe treatment. LEs (white arrowhead), LYS (yellow arrowhead). Representative stills of de-quenching of OG fluorescence in a DQ*hi* LYS **(C)** or DQ*lo* LE **(D).** OG fluorescence in **(E)** DQ*hi* LYS and **(F)** was quantified at t=-2 min and t=3 min after DMSO (black dots) or LLOMe (pink dots) and compared using paired Wilcoxon-Test. (n=15 movies per condition, 3 independent experiments, n.s. = non-significant, ***p<0.001, ****p<0.0001).

### pH gradient collapse is sufficient to phenocopy many LLOMe-driven Rab7 phenotypes

Since LLOMe neutralizes the luminal pH of both LE and LYS (Fig. S1A-C, Fig. 5), we next wondered whether pH collapse alone is sufficient to lead to the observed phenotypic changes in Rab7. To do so, we perturbed endosomal/lysosomal pH pharmacologically (Fig. 6A) and performed Rab7 immunostaining. We first used the weak base ammonium chloride (NH_4_Cl) to neutralize luminal pH (Fig. 6B). Treatment with NH_4_Cl resulted in Rab7 compartments that were enlarged, reduced in number, and greater in Rab7 intensity (Fig. 6B-E). We also directly inhibited the endosomal V-ATPase using Bafilomycin-A1 (BafA1). BafA1 blocks key interactions within the V-ATPase V_0_ domain preventing proton translocation (Wang et al., 2021). Importantly, we used treatment parameters of BafA1 previously shown to have primary effects on endosomal/lysosomal acidification but not secondary effects on endo-lysosome fusion (Klionsky et al., 2008). Rab7+ compartments in cells treated with BafA1 also increased in size, reduced in number, and increased in Rab7 intensity (Fig. 6F-I). To determine if these pH-disruptive treatments caused LMP, we screened for ALIX and Gal3 recruitment in NH_4_Cl- and BafA1-treated cells by fixed immunostaining. Importantly, only LLOMe treatment led to accumulation of these markers (Fig. S6A). Ultrastructural analysis by TEM revealed the appearance of enlarged compartments after LLOMe, NH_4_Cl and BafA1 treatment (Fig. S6B-F), including both lucent and dense enlarged compartments in all three conditions, as previously seen in LLOMe-treated NRK cells (Fig. 1G). Together these data suggest that pH neutralization alone is sufficient to alter Rab7+ compartment intensity, area, and number, but not sufficient to trigger membrane damage.

**Figure 6.**
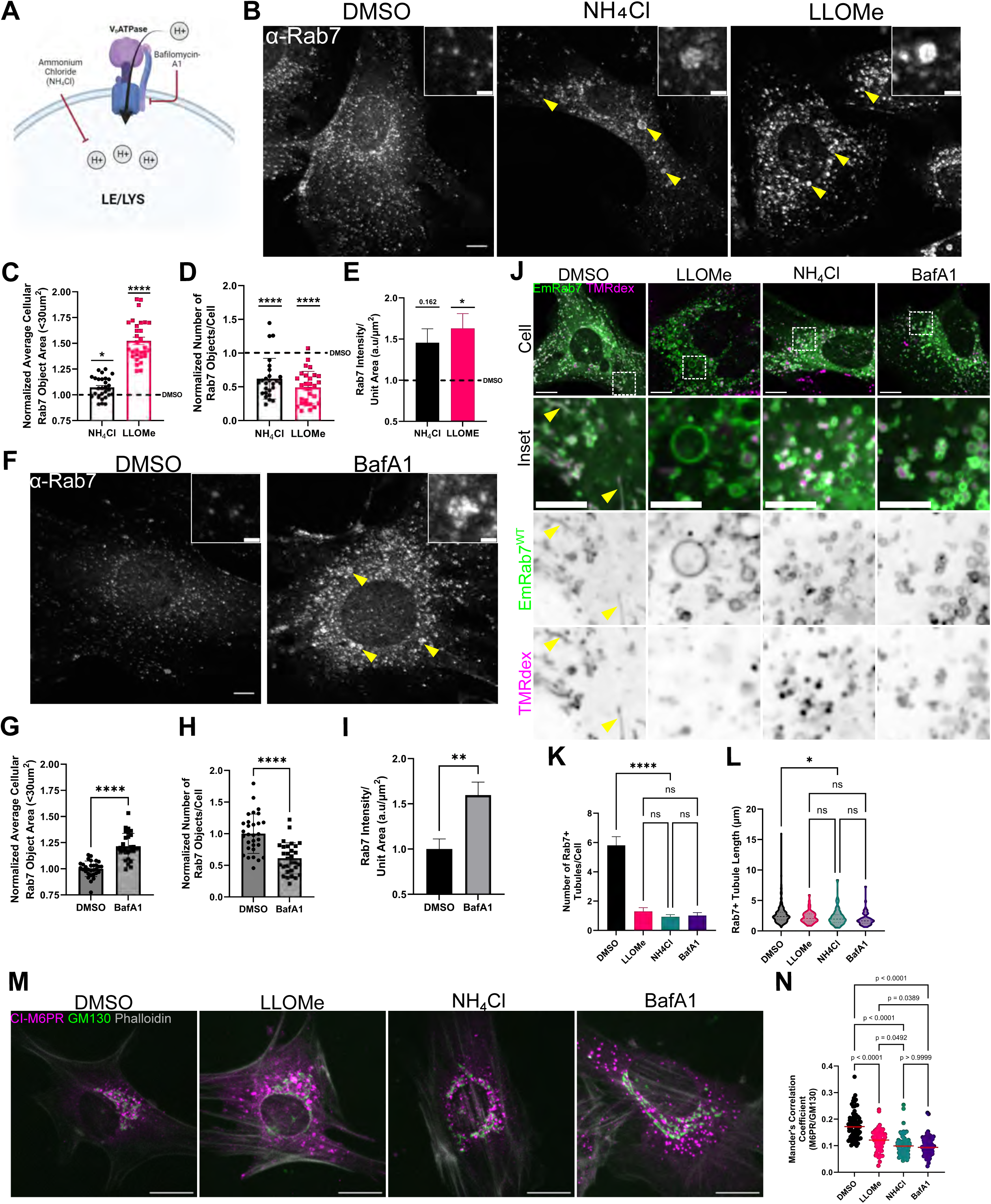
pH gradient collapse is sufficient to phenocopy select LLOMe-driven Rab7 phenotypes. **(A)** LE/LYS pH manipulation with neutralizing base NH_4_Cl and V-ATPase inhibitor Bafilomycin-A1 (BafA1). **(B)** Widefield micrographs of primary MEFs treated with DMSO (1hr), NH_4_Cl (20mM, 1hr), and LLOMe (1mM, 1hr) and immunostained for endogenous Rab7 (scale = 10µm, inset= 2µm). **(C-E)** Using Imaris, Rab7 surfaces were created and filtered for those <30µm^2^ in area. Cumulative cellular surface properties were analyzed for average area of Rab7 surfaces (mean±SEM) **(C)**, number of Rab7 surfaces (mean±SD) **(D)**, and Rab7 intensity per unit area of created surfaces (mean±SEM) **(E)**, per cell. (C-E) were compared using One-Way ANOVA, Welch’s ANOVA, and Kruskal-Wallis, respectively (n=30 FOV, 3 independent experiments) **(F)** Widefield micrographs of primary MEFs treated with DMSO (1hr) and BafA1 (200nM, 2hr) and immunostained for endogenous Rab7 (scale = 10µm, inset= 2µm). **(G-I)** Quantification and comparisons were performed in parallel to those in (C-E). **(J)** Confocal micrographs of Em-Rab7^WT^ transfected primary MEFs loaded with 100µg/mL TMRdex (magenta) for 2hr, prior to DMSO, 1mM LLOMe or 20mM NH_4_Cl for 45mins, or 200nM BafA1 for 1hr. Em-Rab7^WT^ tubules (green) = yellow arrowheads (scale= 10µm, inset= 5µm). **(K)** Quantification of the number of Em-Rab7^WT^+TMRdex+ tubules in (J), compared using Kruskal-Wallis (n=57 (DMSO), 50 (LLOMe), 57 (NH4Cl) and 60 (BafA1) cells, shown mean±SEM, 3 independent experiments). DMSO and LLOMe conditions are the same as Fig. 3 F-H. **(L)** Length quantification of Em-Rab7^WT^+TMRdex+ tubules in (F), compared using Kruskal-Wallis (n=331 (DMSO), 64 (LLOMe), 53 (NH4Cl), and 61 (BafA1) tubules, 3 independent experiments). **(M)** Widefield micrographs of primary MEFs treated with pH drugs as in (B) and (F) and immunostained for endogenous CI-M6PR (magenta) and GM130 (green) (scale =20µm) **(N)** Manders correlation coefficients for M6PR on GM130 space were determined using Imaris and compared using Kruskal-Wallis (n=74 (DMSO), 57 (LLOMe), 69 (NH4Cl), and 63 (BafA1) cells from 2 independent experiments, line = median) (n.s. = not significant, *p<0.05, **p<0.01, ***p<0.001, ****p<0.0001)

We next asked whether direct pH collapse is sufficient to cause endosomal tubulation deficits and CI-M6PR mis-trafficking akin to LLOMe-mediated stress. Assessment of tubulation activity in Em-Rab7^WT^ expressing MEFs loaded with TMR-dextran and treated with NH_4_Cl or BafA1 reduces tubule number and length, as compared to DMSO (Fig. 6J-L). Similarly, endogenous CI-M6PR was mis-localized away from the cis-Golgi marker GM130 in NH_4_Cl- and BafA1-treated COS-7 cells (Fig. 6M-N). Together, these data suggest that pH neutralization is sufficient to disrupt Rab7+ endosome tubulation dynamics and to cause mis-localization of CI-M6PR.

### LLOMe-mediated pH collapse recruits V-ATPase V_1_ subunits to late endosomes and lysosomes independently of Rab7

We have shown that LLOMe-mediated luminal pH neutralization and Rab7 hyper-activation are both sufficient to lead to changes in Rab7+ endosome/lysosomes (including defects in CI-M6PR trafficking and tubulation dynamics). We next sought to address how luminal pH changes, such as from LLOMe treatment, might be connected to the altered activity of cytosolic Rab7. V_0_ subunits of the endosomal V-ATPase are responsive to luminal pH changes (Hurtado-Lorenzo et al., 2006; Marshansky, 2007), making the V-ATPase itself a likely candidate to transmit pH information across the endosomal membrane. In yeast and HEK293T cells, vacuolar and lysosomal pH neutralization increases pump assembly (Shao and Forgac, 2004; Stransky and Forgac, 2015). We thus hypothesized that LLOMe would lead to increased V-ATPase assembly (and thus activity) on endosomes in an attempt to re-establish low luminal pH. To test this, we overexpressed an HA-tagged version of V_1_G_1_ in COS-7 cells and co-immunostained for endogenous Rab7 and the HA-tag. 30 minutes of 1mM LLOMe treatment led to increased V_1_G_1_ on Rab7 endosomes as compared to controls, consistent with increased V-ATPase assembly (Fig. 7A-B). To then test whether V_1_G_1_ recruitment occurred on LEs or LYS, we loaded HA-V_1_G_1_ transfected COS-7 cells with DQ-BSA to mark LYS. After LLOMe treatment, we observed increased recruitment of HA-V_1_G_1_ to both LEs (Dq*lo*; white arrowhead) and LYS (Dq*hi*; yellow arrowhead) Rab7+ compartments (Fig. 7C). These data suggest that LLOMe increased pump assembly on both LEs and LYS. Given that LEs themselves are not extensively damaged (Fig. 4), these data also suggest that membrane damage is not a pre-requisite for V-ATPase assembly in LLOMe.

**Figure 7.**
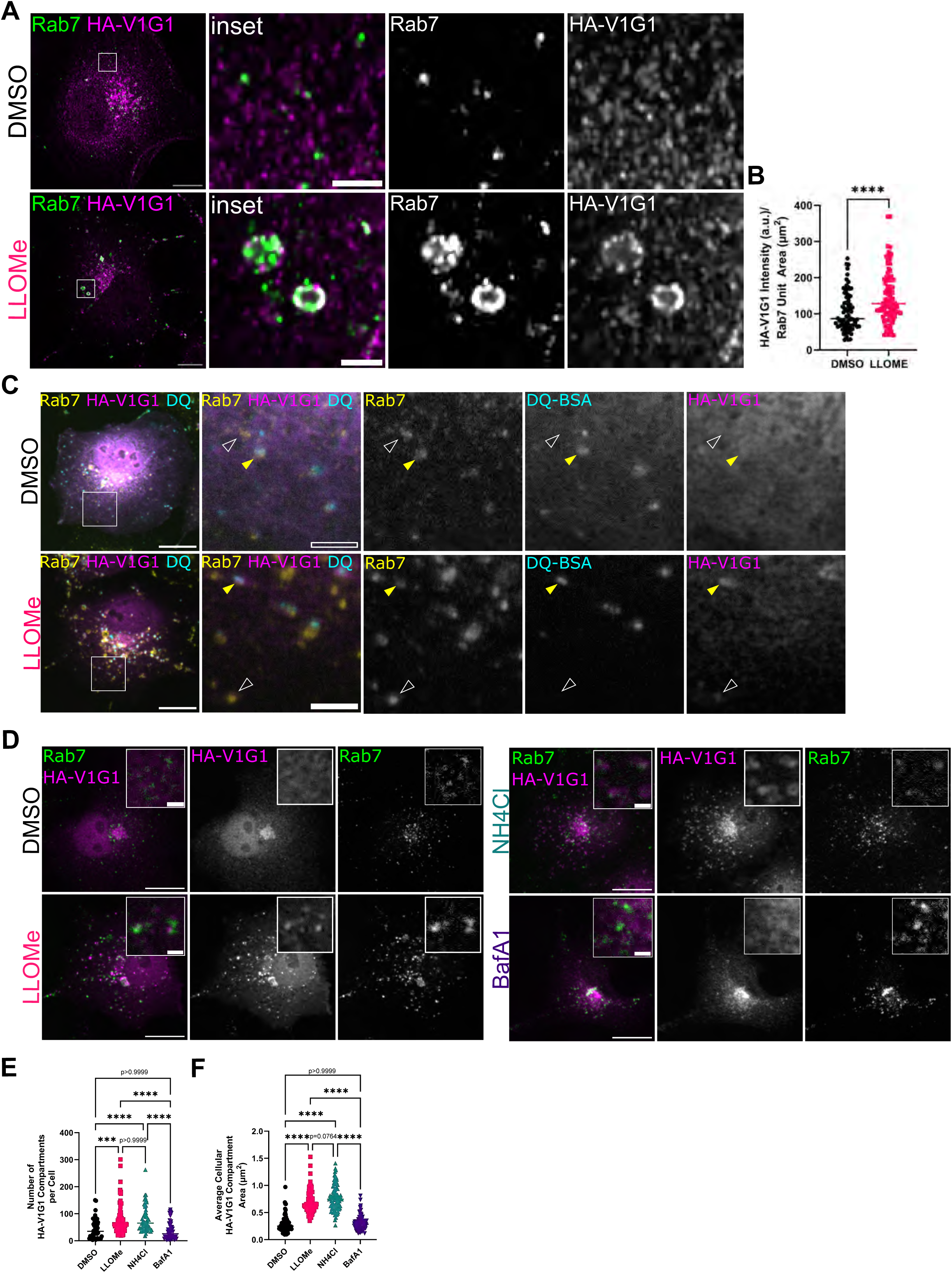
LLOMe-mediated pH collapse recruits the V_1_G_1_ subunit of the V-ATPase to late endosomes and lysosomes. **(A)** Confocal micrographs of HA-V_1_G_1_ transfected COS-7 treated with DMSO or LLOMe (1mM, 30mins), then immunostained for endogenous Rab7 and HA-epitope (scale= 10µm, inset = 2µm). **(B)** Quantification of (A) compared using Mann-Whitney-U (n=76 (DMSO) and 85 (LLOMe) cells, 3 independent experiments, line = median). **(C)** Widefield micrographs of HA-V_1_G_1_ transfected COS-7 loaded with 5µg/mL DQ-BSA (cyan) overnight and treated with DMSO or LLOMe (1mM, 30mins), fixed, and immunostained for endogenous Rab7 (yellow) and HA-epitope (magenta) (scale= 20µm, inset= 5µm). LEs (white arrowhead), LYS (yellow arrowhead) **(D)** Widefield micrographs of HA-V_1_G_1_ transfected COS-7 treated with pH drugs as in Fig. 6B,F, immunostained for endogenous Rab7 (green) and the HA-epitope (magenta) (scale= 20µm inset= 2µm). The **(E)** number and **(F)** area of HA-V_1_G_1_ compartments were quantified and compared using Kruskal-Wallis [n=47 (DMSO), 63 (LLOMe), 56 (NH_4_Cl), 51 (BafA1) HA-V_1_G_1_ transfected cells, 3 independent experiments, line = median] (n.s. = not significant, *p<0.05, **p<0.01, ***p<0.001, ****p<0.0001).

Rab7 indirectly associates with the V-ATPase via a ternary Rab7-RILP-V_1_G_1_ complex (De Luca et al., 2014). Since both Rab7 and V_1_G_1_ levels are increased on endo-lysosomes following LLOMe, we hypothesized that Rab7 activity might control V_1_G_1_ localization and thus V-ATPase activity following LLOMe. To investigate this hypothesis, we overexpressed either GFP-Rab7^WT^, GFP-CA-Rab7^Q67L^, or dominant negative (GFP-DN-Rab7^T22N^) with HA-V_1_G_1_ in COS-7 cells and then treated with LLOMe (Fig. S7A-C). If hyperactivated Rab7 is responsible for HA-V_1_G_1_ assembly on endosomal membranes, we expect GFP-CA-Rab7^Q67L^ to drive HA-V_1_G_1_ assembly regardless of LLOMe treatment, and GFP-DN-Rab7^T22N^ to prevent HA-V_1_G_1_ assembly in LLOMe. Surprisingly, HA-V_1_G_1_ assembly was not increased with GFP-CA-Rab7^Q67L^ expression alone, even though Rab7 is highly recruited and endosomes were enlarged (Fig. S7B,D-E). Similarly, HA-V_1_G_1_ assembly in LLOMe was not inhibited by GFP-DN-Rab7^T22N^ (Fig. S7C-E). Interestingly, GFP-DN-Rab7^T22N^ was mildly recruited to HA-V_1_G_1_ compartments in LLOMe, raising the possibility of a nucleotide-independent mechanism for the initial Rab7 recruitment under these conditions. To test whether Rab7 was required for HA-V_1_G_1_ localization at all, we transiently transfected Cre-GFP and HA-V_1_G_1_ into primary MEFs from Rab7^fl/fl^ mice. Despite the lack of Rab7 in Cre-GFP+ cells, HA-V_1_G_1_ localized normally to endosomes under untreated conditions (Fig. S7F). Together, these data suggest that LLOMe pH neutralization leads to proton pump assembly (V_1_G_1_ recruitment) on both LEs and LYS independently of Rab7.

### V-ATPase assembly contributes to Rab7 membrane stabilization

Since Rab7 was not required for V_1_G_1_ recruitment, we wondered if V-ATPase assembly instead stabilized Rab7, which would constitute a “lumen-to-cytosol”-driven Rab7 regulation mechanism. To begin to support this possibility, we first tested whether Rab7 localized to endosomes/lysosomes in conditions were V_1_ is known to be highly recruited for active acidification, such as starvation, in the absence of pH neutralization. HA-V_1_G_1_ transfected COS-7 cells were exposed to either 24-hour nutrient deprivation in EBSS or 4-hour treatments with Torin-1. These treatments were capable of inducing formation of RFP-LC3 puncta (Fig. S8A) and led to significant changes in V_1_G_1_ compartment number and area (Fig. S8B-C). Notably, endogenous Rab7 that localized to V_1_G_1_ space was also significantly increased (Fig. S8D). Conversely, only Torin-1 led to significant changes in Rab7 puncta number and area (Fig. S8E), and showed V_1_G_1_ enrichment on Rab7 space (Fig. S8F). Together, these data support that active V_1_ recruitment to endosomes is correlated with increased endosomal Rab7, irrespective of the pH at the time of recruitment.

We then determined whether pH neutralization alone was sufficient to recruit HA-V_1_G_1_ to endosomes. Whereas LLOMe and NH_4_Cl treatment led to high degrees of recruitment of HA-V_1_G_1_ to endosomal compartments, BafA1 did not (Fig. 7D-F), despite BafA1 causing pH collapse in COS-7 cells and all other cell types (Fig. S7G-I), consistent with previous literature (Hooper et al., 2022). Therefore, LLOMe and NH_4_Cl lead to pH neutralization with pump assembly (i.e., V_1_G_1_ recruitment) whereas BafA1 leads to pH neutralization without significant pump assembly.

Leveraging differential proton pump assembly in pH-neutralizing conditions, we next asked if Rab7 membrane stability was changed as a result of assembly, speaking to V-ATPase mediated changes to Rab7. We carried out FRAP on Rab7+ endosome/lysosomes in stable Em-Rab7^WT^ NRK cells under various pH-neutralizing conditions and compared them to previously established DMSO and LLOMe conditions (Fig. 3C-E; Fig. 8A, Video 7). Em-Rab7 recovery in NH_4_Cl-treated cells was significantly slower than controls (Fig. 8A-C, teal), indicating increased Rab7 stability on endosomes in both LLOMe and NH_4_Cl. Interestingly, while BafA1-treated Em-Rab7 endosome/lysosomes also recovered significantly slower than controls, they recovered significantly faster and to a greater extent compared to LLOMe (Fig. 8A-C, purple), indicating lower degrees of Rab7 membrane stabilization. Importantly, the randomly selected cells used for bleaching were similar in normalized fluorescence intensity of Em-Rab7^WT^ across conditions (Fig. 8D). Single exponential phase fitting of FRAP curves (Fig. 8I) confirmed these findings and revealed altered half-times (t_1/2_) of recovery following pH neutralizing conditions. Specifically, while the half time in control cells was quick (t_1/2_= 54.87s), LLOMe greatly extended this (t_1/2_= 188.2s), whereas BafA1 was only partially impacted (t_1/2_= 66.1s). Putting these data together, pH neutralization conditions that enhanced V-ATPase assembly (i.e., LLOMe and NH_4_Cl) stabilized Rab7 association with LE/LYS membranes, whereas conditions without V-ATPase assembly (i.e., BafA1) did so to a significantly lesser degree (Fig. 8J).

**Figure 8.**
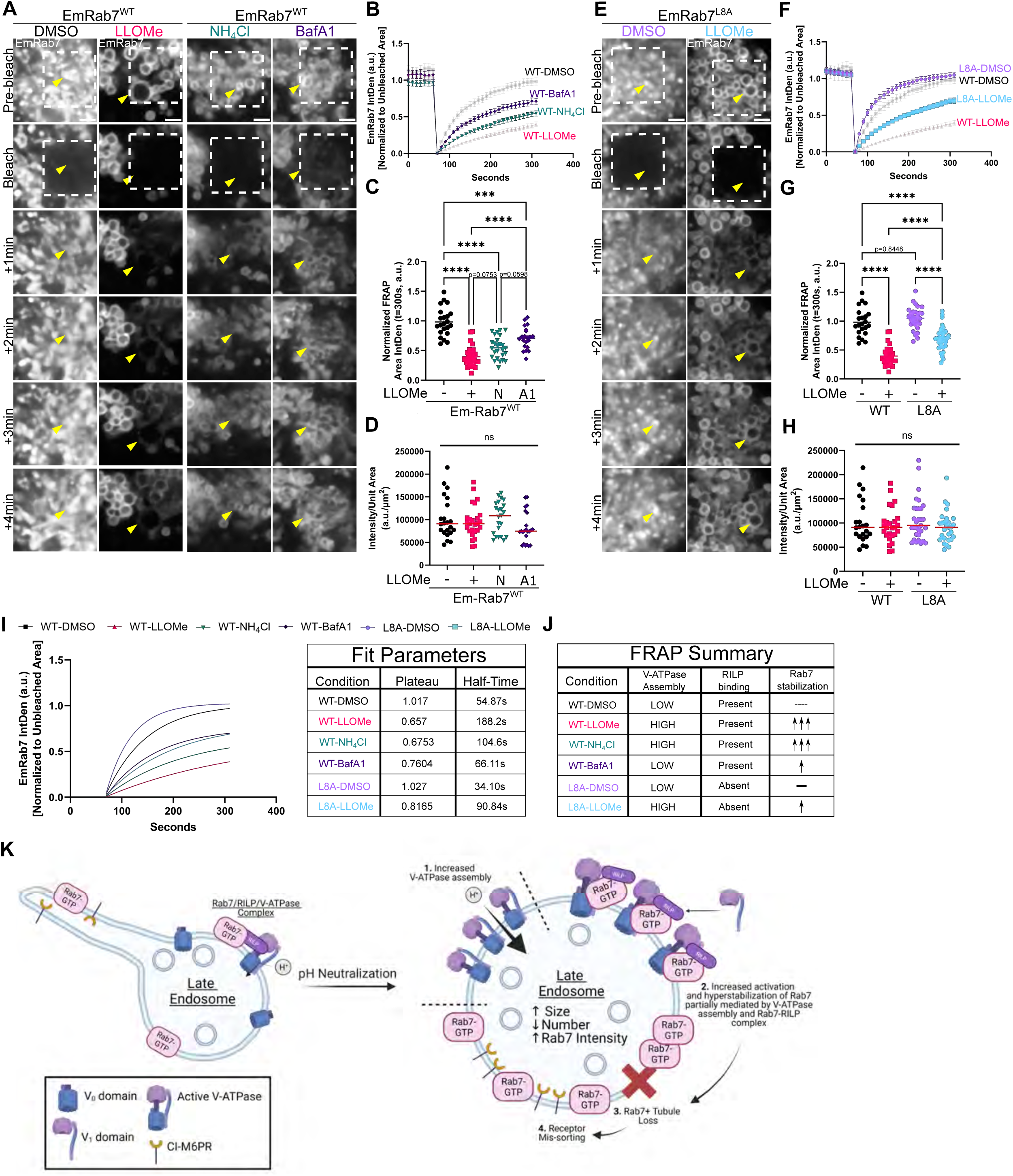
LLOMe drives membrane stabilization and hyperactivation of Rab7 through influence from V-ATPase assembly and RILP. **(A)** Representative stills from confocal FRAP live imaging (Video 7) of Em-Rab7^WT^ NRK cells under DMSO (10mins, n=23), LLOMe (1mM, 10mins, n=27), NH_4_Cl (20mM, 10mins, n=26), or BafA1 (200nM, 1hr, n=21) conditions. Bleached area indicated by white box (33.0625µm^2^) with representative Rab7+ endosomes= yellow arrowhead (scale = 2µm). **(B)** FRAP recovery over time from (A). **(C)** Mean FRAP area intensity values at t= 300sec (Y_max_) and compared using One-Way ANOVA (n= as in (A), 3 independent experiments, line = median, ***p<0.001, ****p<0.0001). DMSO and LLOMe conditions are from Fig. 3 C-E. **(D)** Mean normalized Em-Rab7 intensity of FRAP cells prior to bleaching. Red bar= median, compared using Kruskal-Wallis (n= as in (A)). **(E)** Representative stills from confocal FRAP live imaging (Video 7) of Em-Rab7^L8A^ NRK cells under DMSO (10mins, n=28) or LLOMe (10mins, n=30). 33.0625µm^2^ bleached area indicated by white box. Representative Rab7+ endosomes = yellow arrowhead. (scale = 2µm). **(F)** FRAP recovery over time in (E). **(G)** Mean FRAP area intensity values at t= 300sec (Y_max_) and compared using One-Way ANOVA (n= as listed in (E), 3 independent experiments, line = median, ***p<0.001, ****p<0.0001). Wild-type data in (E-H) are identical to that of (A-D); mutant data has been separated for clarity. **(H)** Mean normalized Em-Rab7 intensity of L8A FRAP cells, prior to bleaching. Red bar= median, compared using Kruskal-Wallis (n= as in (E)). **(I) *Left:*** One-way exponential phase association fitted curves for all conditions following bleach. ***Right:*** Parameters from fitted curves indicating the plateau values and half-time for each condition. **(J)** FRAP summary table for all conditions based on V-ATPase assembly and RILP binding. **(K)** Our model for pH regulation on Rab7-positive late endosomes proposes that upon pH neutralization there is increased V-ATPase assembly (1.), via a RILP-V-ATPase axis, stabilizing Rab7 in a hyperactive form (2.) which contributes to the loss of endosomal tubulation (3.) and proper CI-M6PR trafficking (4.).

Rab7 indirectly associates with the V-ATPase via a ternary Rab7-RILP-V_1_G_1_ complex. We thus investigated if Rab7 stabilization on membranes in response to LLOMe required this complex be intact. Since knockdown of RILP is prone to off-target effects (Yap et al., 2023), we used a point mutant in Rab7 (Em-Rab7^L8A^) which cannot bind RILP (Wu et al., 2005) and thus lacks V_1_G_1_ binding. We created a stable NRK line overexpressing mutant Em-Rab7^L8A^. Normalized fluorescence intensity of Em-Rab7 signal in WT and L8A groups was similar in selected cells (Fig. 8H). Given Rab7 is non-essential for HA-V_1_G_1_ responses to LLOMe (Fig. S7), V-ATPase assembly is intact under these conditions. We then subjected these cells to FRAP experiments. In DMSO, Em-Rab7^L8A^ recovered similarly to Em-Rab7^WT^ following photobleaching (Fig. 8E-G, light purple). However, in LLOMe, recovery of Em-Rab7^L8A^ was significantly faster than that of Em-Rab7^WT^ (Fig. 8E-G, blue). Further, plateau values of LLOMe-treated Em-Rab7^L8A^ recovered to a significantly greater degree than LLOMe-treated Em-Rab7^WT^ cells (Fig. 8G). Single exponential fitting of FRAP curves (Fig. 8I) revealed the half time in LLOMe treated Em-Rab7^L8A^ was much shorter than the equivalent treatment in Em-Rab7^WT^ cells (t_1/2_= 90.84s vs. 188.2s). Interestingly, the fitted recovery of Em-Rab7^L8A^ in LLOMe closely resembled that of BafA1 treatment (Fig. 8I). Since Rab7 can dimerize, we next tested whether depletion of endogenous Rab7^WT^ would lead to even greater FRAP recovery of Em-Rab7^L8A^. We depleted endogenous Rab7 in NRK using an siRNA that we previously validated (Yap et al., 2022). We found a 50% decrease in endogenous Rab7 levels after 4 days of knockdown, in concordance with our previous validation (Fig. S8G-H). This 50% depletion in stable EmRab7^L8A^ mutant NRK cells did not significantly impact FRAP recovery in either DMSO or LLOMe conditions, as compared to a non-targeting control, suggesting the endogenous Rab7 pool is not a significant player in determining recovery of Em-Rab7^L8A^ (Fig. S8I-K). These data together suggest that Rab7 is less stabilized on membranes following LLOMe in the absence of RILP binding (Fig. 8J). Notably, Em-Rab7^L8A^ compartments were still vacuolar and partially stabilized Rab7 in LLOMe, suggesting that other mechanisms exist beyond RILP-mediated stabilization. Since both BafA1 and Em-Rab7^L8A^ conditions independently led to partial rescue of Em-Rab7 recovery relative to LLOMe, we suggest that together V-ATPase assembly and RILP contribute to Rab7 membrane stabilization in pH-neutralizing conditions.

## Discussion

In summary, the present study finds that the behavior of Rab7+ late endosomes upstream of damaged lysosomes is profoundly altered by pH neutralization: increased V-ATPase assembly (Step 1, Fig. 8K) promotes activation and membrane stabilization of Rab7 (Step 2, Fig. 8K), leading to decreased membrane tubulation of endosomes (Step 3, Fig. 8K) and faulty trafficking of CI-M6PR (Step 4, Fig. 8K). In a previous report, chloroquine- or sirasamine-driven LMP damaged 30% of Rab7+ compartments. This population includes LYS and LEs/EEs since they were not distinguished from each other (Du Rietz et al., 2020). Notably, our work carefully defined compartmental markers and confirmed that the majority of Rab7+ compartments with membrane permeabilization were LYS, whereas Rab7+ LEs rapidly respond to LLOMe with pH neutralization but do not exhibit LMP. The acute LE response is characterized by rapid enlargement from pH neutralization, which led to increased V-ATPase assembly on both undamaged LEs and damaged LYS. Whereas pH neutralization due to vacuolar damage in *Salmonella* infection (Xu et al., 2019) or LLOMe treatment (Cross et al., 2023) assembles V-ATPase for autophagy conjugation, our data reveal the novel notion that V-ATPase assembly in sterile (non-infection) pH-neutralized late endosomes leads to increased activation and membrane stability of Rab7, which we found to be further influenced by the ability of Rab7 to bind its effector RILP.

### The relationship between V-ATPase, Rab7 and endosomal pH

The current assumption is that active Rab7 contributes to acidification of endosomes as they mature into functional lysosomes based on a correlation between lower pH and increased Rab7 on endosomes, and early data suggesting DN-Rab7^T22N^ affects lysosomal pH (Bucci et al., 2000). This notion is supported by data including RILP controlling V_1_G_1_ stability, increased pH after RILP depletion, and the correlation of lower levels of Rab7 on less acidic peripheral lysosomes (De Luca et al., 2014; Johnson et al., 2016). However, definitive data demonstrating the ability of active Rab7 to recruit pump subunits or tune luminal pH is lacking. More recent Lysotracker data in Rab7-KO MEFs and MDCK cells demonstrate Rab7 is dispensable for maintenance of endosomal pH (Kuchitsu et al., 2018; Liang et al., 2023). Now, we further demonstrate that Rab7 is dispensable for the assembly of V_1_G_1_ on endosomes, given complete knockout of Rab7 could not deplete endosomal association of V_1_G_1_. Collectively, these data point to the importance of RILP in acidification and stabilization of Rab7.

Whereas initial work suggested that Rab7 was upstream of V-ATPase and regulated acidification via V_1_G_1_ (De Luca et al., 2014), newer work suggested instead that Rab7 membrane recruitment and stabilization may be downstream of the V-ATPase subunit V_0_a_3_. Knockout of V_0_a_3_ eliminates Rab7 association with lysosomes and anterograde trafficking (Matsumoto et al., 2018). BafA1 also decreases anterograde lysosomal trafficking and depletes constitutively active Rab7 in complex with pump subunits (Matsumoto and Nakanishi-Matsui, 2019). Whereas these data identify the V_0_a_3_ subunit, our data more generally implicate an assembled V-ATPase in contributing to the control of Rab7 activation state. In scenarios with pH neutralization and extensive pump assembly (i.e., LLOMe or NH_4_Cl), we observed high degrees of membrane stabilization of Rab7 by FRAP. On the contrary, low pump assembly conditions (i.e., BafA1) had significantly destabilized Rab7 on endosomal membranes.

In our interpretation, these data provide evidence that Rab7 activity is subject to regulation by luminal pH-driven V-ATPase assembly when pH is too alkaline. This might include, for example, the maturational transition from more alkaline EE to acidic LEs. Our Rab7-mutant and knockout data suggest that physiologic acidification is Rab7-independent, and rather V-ATPase reciprocally increases the presence of active Rab7 on more mature compartments. In support of the notion, we demonstrate that physiologic starvation conditions, such as in EBSS or direct mTOR inhibition, increased Rab7 on V_1_G_1_ membranes, when acidification occurs without neutralization (Collins and Forgac, 2020; McGuire and Forgac, 2018; Ratto et al., 2022; Stransky and Forgac, 2015). Our data and others also support a role for pump-mediated initiation of Rab7 activation. Previous data suggests V-ATPase may contribute to recruitment of inactive Rab7 and its cognate GEF to the membrane (Matsumoto et al., 2022). Our data further support this possibility, as DN-Rab7^T22N^ is modestly recruited to V_1_G_1_ positive membranes following LLOMe.

Together, it appears that fine tuning endosomal pH involves extensive communication between the V-ATPase and Rab7. Our new data and others are consistent with the idea that V-ATPase is a main driver of Rab7 activation and stabilization on membranes, which is akin to similarly described V-ATPase driven recruitment of the ARF-GEF ARNO for Arf6 activation (Maranda et al., 2001). Our results are further relevant to all instances of endosomal/lysosomal neutralization, including the use of lysosomotropic pH-neutralizing therapeutics, such as selective-serotonin reuptake inhibitors (SSRIs) or atypical antipsychotic therapeutics (Kuzu et al., 2017; Kuwahara et al., 2020; Lakpa et al., 2021), which may similarly cause pH neutralization in LEs and disruption of membrane trafficking in the absence of LMP.

### Endosomal membrane stabilization of active Rab7 is influenced by V-ATPase assembly and RILP binding: an effector balance?

Modest changes in Rab7 activation as a result of pump assembly, downstream of pH sensing, could have profound effects on effector binding important for control of endosomal transitions and cellular membrane trafficking. As evidence of this, the extreme cases of pH neutralization we explore in this paper disrupt normal Rab7-positive tubulation and CI-M6PR trafficking events. These tubulation events are thought to be dependent on dynein-RILP effector complexes (Mrakovic et al., 2012). In our FRAP data, we demonstrate that the increased membrane stabilization of Rab7 is partially due to Rab7 interactions with RILP, as Rab7^L8A^ mutants recover quicker than Rab7^WT^. This is consistent with previous data from both our lab and others that demonstrate overexpressed RILP massively stabilized Rab7 on membranes (Jordens et al., 2001; Yap et al., 2022). Collectively, these data begin to suggest a complex balance of Rab7 effectors on late endosomes that depend on RILP as a limiting factor. We speculate that increased V_1_-RILP-Rab7 interactions in pH neutralization responses sequesters active Rab7 away from other effectors which rely on RILP interactions. CI-M6PR mis-sorting and tubulation deficits might thus be a consequence of a disrupted Rab7 effector balance. Rab7 effector balance is an exciting avenue of future exploration, as it may play key roles in long range trafficking and degradation of important trophic molecules relevant to disease (Mulligan and Winckler, 2023).

### Rab7 tubulation and lysophagy

Our data demonstrate a drastic change in the tubulation activity of Rab7+ compartments after LLOMe. In contrast to the appearance of LAMP1-negative tubules on LAMP1-positive LYS after extensive damage (>1h LLOMe) (Bonet-Ponce et al. 2020) or tubules formed during LMP recovery (Bhattacharya et al. 2023), we observe an acute loss of Rab7+ tubulation in a short temporal window following LLOMe exposure. Recovery of lysosomal tubules from LLOMe requires Rab7 inactivation by the GAP TBC1D15 (Bhattacharya et al., 2023). We speculate that LLOMe initially disrupts normal Rab7-positive tubulation via hyperactivation, which affects the retrieval of membrane cargos that should not enter lysosomes (e.g. CI-M6PR), whereas Rab7-negative tubulation events with separate functions may still occur at later stages of LMP (Bonet-Ponce et al. 2020; Bhattacharya et al. 2023). We further suggest that the rapid loss of tubulation in LLOMe partially contributes to the Rab7 compartment phenotypes (e.g., size) observed in LLOMe. Other factors such as excessive fusion events or lipid exchange mechanisms might play additional roles (Radulovic et al., 2022; Tan and Finkel, 2022).

Finally, V-ATPase assembly-driven Rab7 activation may serve as an accelerant of the endo-lysosomal system. As V-ATPase assembly and acidification were recently linked to increased degradation in previously quiescent compartments (Ratto et al., 2022), it is possible that this mechanism serves to activate a pool of initially weakly-degradative LEs to generate functional lysosomes in order to maintain proteostasis or progress to lysophagy. Given that in other contexts Rab7 activation provides control over the selective autophagy of organelles (Jimenez-Orgaz et al., 2018), future studies are needed to elucidate the definitive downstream role of undamaged, actively acidifying LEs following lysosomal disruption for lysophagic resolution.

Collectively, our data further raise new questions about Rab7 effector selection and its connection to luminal pH, with considerations for lysophagy progression. This mechanistic link likely underlies dynamic changes in homeostatic endosomal behavior in health and might be disrupted in several genetic disorders, including neurodegenerative diseases (Colacurcio and Nixon, 2016; Pescosolido et al., 2021). This study therefore broadens our understanding of physiologic pH regulation, control of Rab7 activity and effector binding, and ultimately cellular responses to LE/LYS disruptions.

## Materials and Methods

### Immortalized cell line culture and transfection

Normal rat kidney (NRK) cells (ATCC CRL6509) and COS-7 cells (UVA Tissue Culture Facility) were maintained in DMEM with glucose and GlutaMAX (Gibco 10566-016) plus 10% fetal bovine serum (FBS) without antibiotics. All cells were incubated at 37°C and maintained at 5% CO_2_. For fixed imaging, cells were plated on coverslips coated with 10 µg/mL fibronectin (Sigma) for 1 hour at 37°C. For live imaging, cells were plated on 35mm dishes with 14mm glass bottom inserts (MatTek) coated with 10µg/mL fibronectin for 1 hour at 37°C. Transfections into NRK and COS-7 cells for fixed or live imaging experiments and for GST-RILP Rab7 activity pulldowns were performed using Lipofectamine 2000 (Invitrogen) per manufacturer’s instructions. Downstream applications were conducted 24 hours post-transfection unless otherwise stated. For establishment of stable Emerald-Rab7-WT and Em-Rab7-L8A NRK cell lines, Em-Rab7^WT^ or Em-Rab7^L8A^ were transfected into NRK cells. 24 hours post transfection, cells were maintained on 100µg/mL geneticin (Gibco) in complete DMEM. After 5 days of selection, individual cell colonies were selected and expanded in geneticin-containing medium. Purity of clones was screened for homogenous Em-Rab7 expression by fluorescence microscopy. Pure clones were expanded, frozen, and maintained in geneticin-containing DMEM for downstream applications.

### Primary mouse embryonic fibroblast isolation and nucleofection

All experiments were performed in accordance with the University of Virginia Institutional Animal Care and Use Committee guidelines and regulations. Primary mouse embryonic fibroblasts (MEFs) were isolated from E13.5 Rab7^fl/fl^ mice (Kawamura et al., 2012) (Riken BioResource Research Center #RBRC05600). Embryos were sterilely isolated from fluid sacs and torsos were isolated by removal of the head, limbs and tail. Internal organs and spinal cord were then removed by forceps. Remaining tissue was triturated in 0.05% trypsin-EDTA then incubated at 37°C for 10 minutes. An equivalent volume of 20% FBS in MEF medium [DMEM with glucose and L-glutamine (Gibco 11965-092)] was then added and the tissue was re-triturated. Cells were then spun, re-suspended, and plated in MEF medium containing 10% FBS for expansion, freezing, or downstream applications. Upon use, MEFs were maintained in MEF medium containing 10% FBS without antibiotics. For single or double plasmid expression experiments (e.g., Em-Rab7 overexpression or Cre-recombinase knockouts), MEFs were nucleofected using Ingenio Solution electroporation reagent (Mirus Bio) and Lonza AMAXA Nucleofector II. Briefly, 1.5 million MEFs and 5µg total DNA were diluted in 100µL of Ingenio Solution in a 0.2cm cuvette (Mirus). MEFs were nucleofected using AMAXA program A-023 (MEF). Cells were immediately recovered in 900µL of pre-warmed complete MEF medium and then diluted and plated on 10µg/mL fibronectin coated coverslips. For Rab7 knockout experiments, downstream analyses were conducted three days post pCAG-Cre-GFP nucleofection. For live imaging use, MEFs were plated on 35mm dishes with 14mm glass bottom inserts (MatTek) coated with 10µg/mL fibronectin for 1 hour at 37°C.

### Plasmids

The plasmids used in this study are as follows:

Emerald-Rab7^WT^ (Addgene #54244, Davidson Lab), GFP-Rab7^WT^ (Addgene #12605, Pagano Lab), GFP-Rab7^Q67L^ (gift from Dr. James Casanova, University of Virginia), GFP-Rab7^T22N^ (Addgene #12660, Pagano Lab), pCAG-Cre-GFP (Addgene #13776, Cepko Lab), pCDNA3-2xHA-V1G1 (gift from Dr. Cecilia Bucci, University of Salento), pET-His-GST-tev-LIC (Addgene #29655, Gradia Lab), pGEX-4T-3-mR7BD (Addgene #79149, Edinger Lab), mCherry (Clontech), pmRFP-LC3 (Addgene #21075, Yoshimori Lab). To generate Emerald-Rab7^L8A^, Rab7a with a point mutation at residue 8 from leucine to alanine (L8A) was cloned into pEmerald-C1 (Addgene#54734, Davidson Lab) at XhoI-BamHI sites by Genscript, using gene synthesis.

### Primary Antibodies

The list of primary antibodies used in this study are as follows: Anti-Rab7 (rabbit monoclonal; 1:100 IF; abcam 137029; RRID:AB_2629474), Anti-Rab7 (rabbit polyclonal; 1:1000 WB; proteintech 55469-1-AP; RRID:AB_11182831), Anti-Cathepsin B (goat polyclonal; 1:200 IF; R&D Systems AF965; RRID: AB_2086949), Anti-Cathepsin D (goat polyclonal; 1:200 IF; R&D Systems AF1029; RRID: AB_2087094), anti-LAMP1 (mouse monoclonal; 1:1000 IF; Enzo Life Sciences ADI-VAM-EN001; RRID:AB_2038958), anti-GM130 (mouse monoclonal; 1:500 IF; BD Biosciences 610822 RRID: AB_398141), anti-TGN38 (mouse monoclonal; 1:200 IF; BD Biosciences 610898, RRID:AB_398215), anti-Vps35 (goat polyclonal; 1:600 IF, 1:2000 WB; abcam 10099; RRID:AB_296841), anti-EEA1 (mouse monoclonal; 1:100 IF; BD Biosciences 610456; RRID: AB_399409), anti-CI-M6PR (rabbit monoclonal; 1:200 IF; abcam 124767; RRID:AB_10974087), anti-ALIX (mouse monoclonal; 1:100 IF; Santa Cruz sc-53540; RRID: AB_673819), anti-Galectin-3 (mouse monoclonal; 1:100 IF; Santa Cruz sc-25279; RRID: AB_627656), anti-Galectin-3 (rat monoclonal; 1:100 IF; Santa Cruz sc-23938; RRID: AB_627658), anti-GFP (rabbit polyclonal; 1:2000 WB; proteintech 50430-2-AP; RRID: AB_11042881), anti-GST (mouse; 1:1000 WB; Santa Cruz sc-138; RRID: AB_627677); anti-HA (mouse monoclonal; 1:100 IF, 1:1000 WB; Santa Cruz sc-7392; RRID: AB_627809); anti-HA (rabbit; 1:100 IF, 1:1000 WB; Santa Cruz sc-805; RRID: AB_631618).

### Secondary Antibodies

For immunocytochemistry, the following Alexa Fluor–coupled antibodies from Invitrogen were used: Alexa405-Phalloidin (A30104), Alexa488 donkey anti-mouse (A21202; RRID: AB_2535792), Alexa488 donkey anti-rabbit (A21206; RRID: AB_ 2534102), Alexa488 donkey anti-goat (A11055; RRID: AB_2534013), Alexa568 donkey anti-mouse (A10037; RRID: AB_2534017), Alexa568 donkey anti-rabbit (A10042, RRID: AB_2534104), Alexa568 donkey anti-goat (A11057; RRID: AB_162546), Alexa647 donkey anti-mouse (A31571; RRID: AB_2536183), Alexa647 donkey anti-rabbit (A31573; RRID: AB_2535864), Alexa647 donkey anti-goat (A21447; RRID: AB_2340373). The following antibodies from Jackson ImmunoResearch Laboratories, Inc. were used: Alexa488 donkey anti-rat (712-545-153; RRID: AB_2340684), Alexa568 donkey anti-rat (712-585-153; RRID: AB_2340689), Alexa647 donkey anti-rat (712-605-153; RRID: AB_2340694), DyLight405 donkey anti-goat (705-475-147; RRID: AB_2340427), DyLight405 donkey anti-rat (712-475-153; RRID: AB_2340681). For western blotting, the following Jackson ImmunoResearch Laboratories, Inc. antibodies were utilized: donkey anti-mouse Alexa680 (715–625-151; RRID: AB_2340869), donkey anti-mouse Alexa800 (715-655-151; RRID: AB_2340871), donkey anti-rabbit Alexa680 (711-625-152; RRID: AB_2340627), donkey anti-rabbit Alexa790 (111-655-144; RRID: AB_2338086).

### Immunocytochemistry and image acquisition

Immunocytochemistry was conducted as described (Mulligan et al., 2023). Briefly, cells were fixed in 4% paraformaldehyde/3% sucrose in PBS at room temperature for 15 minutes. Cells were then blocked and permeabilized using PBS/5% normal donkey serum/0.2% TritonX-100 or 0.1% saponin for 20 minutes at room temperature. Saponin permeabilization was used when staining for lysosomal membrane proteins (e.g., LAMP1, LAMP2); TritonX-100 was used for all others. Primary and secondary antibodies were diluted in 1% bovine serum albumin (BSA)/PBS and incubated for 1 hour and 30 minutes, respectively, at room temperature. 0.1% saponin was included in BSA antibody dilutions when staining for lysosomal membrane proteins. Coverslips were mounted in ProLong Gold antifade mounting medium (Invitrogen).

For widefield imaging, mounted coverslips were viewed using a Zeiss Z1-Observer with a 40x objective (enhanced chemiluminescence Plan-Neofluor 40x/0.9 Pol working distance = 0.41). Apotome structured illumination was used for all images. Images were captured with the Axiocam 503 camera using Zen software (2012 Blue edition, Zeiss) and identically processed in FIJI software. On average, 10 representative fields per condition per experiment were captured, for 3 independent experiments.

For live-cell spinning disk confocal microscopy, medium in live cell imaging dishes was replaced with phenol-red free DMEM (Gibco 31053-028) plus 10% FBS and 1X GlutaMAX (Gibco 35050-061) prior to imaging. Cells were viewed using a temperature-controlled (37°C) Nikon TE 2000 microscope using a 60X/NA1.45 oil objective, equipped with Yokogawa CSU 10 spinning disc and images were captured with a 512X512 Hamamatsu 9100c-13 EM-BT camera. On average, multicolor images were collected at a rate of 1 image/3 seconds for indicated times, with perfect focus on, up to 30 minutes (600 frames) and compiled using FIJI software. Vehicle (DMSO) or LLOMe (final concentration 1mM) were added in a pre-warmed bolus at 2 minutes (frame 40) live on the stage for all live-imaging experiments. On average, 10-15 movies per condition were collected from 3 independent experiments.

For traditional confocal microscopy, cells plated in live imaging dishes with phenol-red free DMEM (Gibco 31053-028) plus 10% FBS and 1X GlutaMAX (Gibco 35050-061) or mounted coverslips were viewed using a Nikon Ti2-E inverted microscope with AX-R confocal microscopy system and 25 mm FOV resonant scanner/25 mm FOV galvano scanner, with an NSPARC detector unit. The system was controlled and images captured using photomultipliers and the Nikon NIS Elements C software. Cells were viewed using a 60X/NA1.40 oil objective and maintained in a temperature-controlled system at 37°C with perfect focus during live image acquisition. Captured images and movies were identically processed using Elements Denoise.ai followed by compiling in FIJI software. On average, 20-30 movies (FRAP), and 45-60 images (tubulation) per condition were collected from 3 independent experiments.

For Airyscan imaging, mounted coverslips were viewed using the Zeiss 980 NLO system using an inverted Axio Observer microscope with a 63X/NA1.40 oil objective and an 8Y-airyscan detector. Images were collected using Zeiss Zen Blue software on Super-Resolution-4Y mode with 0.13µm z-sections, and z-stacks were identically processed using Zen Airyscan 3-D Processing. On average, 3 representative fields per condition were captured, for 3 independent experiments (approximately 30 total cells per condition).

### Lysosomal perturbation treatments

For induction of lysosomal damage, all cells were incubated in complete DMEM or MEF medium with 10% FBS, supplemented with DMSO (vehicle, 0.33%v/v, Millipore Sigma) or L-leucyl-L-leucine dimethyl ester (LLOMe, Sigma) at 1mM (unless otherwise noted) for times indicated. DMSO and LLOMe were equilibrated in fresh, pre-warmed (37°C) complete DMEM with 10% FBS prior to addition to cells.

All other treatments were carried out in complete DMEM or MEF medium with 10% FBS. Bafilomycin-A1 (Sigma) was dissolved in DMSO used at a final concentration of 200nM for 2 hours. Ammonium chloride (Fisher) was dissolved in sterile water and used at a final concentration of 20mM for 1 hour. ML-SA1 (Tocris) was dissolved in DMSO and used at a final concentration of 20µM for 5-30 minutes. For calcium chelation experiments, cells were pre-treated with 10µM BAPTA-AM (Tocris) for 30 minutes. Then, cells were treated with fresh medium containing either DMSO or LLOMe in the presence of an additional 10µM BAPTA-AM for indicated times.

### Electron Microscopy

NRK cells or MEFs were grown on 1% fibronectin (in PBS), and 0.1 mg/mL poly-L-lysine (in ddH2O) coated 6mm sapphire disks (Technotrade International) in complete DMEM, or complete MEF medium, at 37°C and treated with 45 minutes of DMSO, 1mM LLOMe, 20mM NH_4_Cl, or 2 hours of 200nM BafA1 before immediate cryo-fixation by high pressure freezing (Leica EM ICE High Pressure Freezer), with 20% BSA in DMEM as a cryoprotectant. Samples were freeze substituted for three days in acetone containing 0.2g/mL methylene green (Spectrum Chemical MFG. GRP.), 0.1% uranyl acetate (Polysciences Inc) and 1% osmium tetroxide (Electron Microscopy Sciences) using Leica EM Automatic Freeze Substitution machine (Leica AFS2). Freeze substituted samples were then infiltrated with increasing Epon/Araldite (Electron Microscopy Sciences) in acetone until they were infiltrated with pure Epon/Araldite, over the course of one day. Epon infiltrated samples were embedded in resin and allowed to set for two days at 60°C followed by mounting on dummy resin blocks. Samples were then sectioned at 70nm using an ultramicrotome (Reichert Ultracuts, Leica) and collected onto 0.7% Pioloform (Ted Pella Inc) coated copper slot grids 2mm x 1mm Ø 3.05mm (Electron Microscopy Sciences). Post staining of samples on grids was performed with 2% uranyl acetate in 70% methanol followed by 0.4% lead citrate in ddH2O. Transmission electron micrographs were collected using a Tecnai F20 Transmission Microscope operated at 200kV and captured using a 4K x 4K UltraScan CCD camera.

### Live Imaging Assays

#### Lysosomal pH Collapse Readouts

NRK, COS-7, or MEF cells were loaded with 100µg/mL10kDa fluorescein (FITC) dextran plus 100µg/mL 70kDa tetramethylrhodamine (TMR) dextran (Invitrogen) in complete DMEM at 37°C for 16 hours. After washing with 1X DPBS, dextrans were chased to lysosomes for an additional 4 hours at 37°C in phenol red-free complete DMEM. Live time lapse images were captured via spinning disk microscopy with LLOMe added on the stage after 2 minutes (Fig S1) or still images were collected following 2-hour treatment courses (Fig. S7). Change in cellular FITC fluorescence over time was plotted relative to initial cellular FITC intensity, relative to TMR (Fig S1) or ratio of FITC:TMR dextran was reported within TMR ROIs (Fig. S7).

#### Em-Rab7 Live Imaging of Late Endosomes versus Lysosomes

Stable Em-Rab7^WT^ NRK cells were loaded with 5µg/mL DQ-BSA (Invitrogen) in complete DMEM at 37°C for 16 hours. The following morning, cells were washed with 1X DPBS and chased for 4 hours in complete DMEM at 37°C. DQ-loaded Em-Rab7^WT^ were then imaged at 37°C with spinning disk confocal microscopy, with DMSO or 1mM LLOMe added live on the stage.

#### pH Collapse of Late Endosomes versus Lysosomes

NRK cells were loaded with 5µg/mL DQ-BSA (Invitrogen) in complete DMEM at 37°C for 16 hours. The following morning, cells were washed with 1X DPBS and then loaded with 100µg/mL10kDa Oregon Green dextran (OGdex) for 2 hours. Following 2-hour OGdex loading, cells were washed with 1X DPBS and replaced with complete phenol-red free DMEM and imaged with spinning disk confocal microscopy. DMSO or 1mM LLOMe was added live on the stage during movie acquisition.

#### Dextran tubulation

For Fig. 3 F,J and Fig. 6 J, MEFs nucleofected with Em-Rab7^WT^ or Em-Rab7^Q67L^ were loaded with 100µg/mL10kDa tetramethylrhodamine (TMR) dextran for 2 hours. After washing with sterile 1X DPBS, the medium was replaced with phenol red-free complete DMEM supplemented with DMSO or 1mM LLOMe for 30 minutes, 20mM NH_4_Cl for 20mins, or 200nM BafA1 for 1 hour at 37°C. Still images of dynamic tubules were then captured by live confocal microscopy (Nikon Ti2-E with AX-R) immediately following recovery.

For Fig. 3 I and Video 4, MEFs were loaded with 5µg/mL DQ-BSA in complete DMEM at 37°C for 16 hours. The following day, after washing with sterile 1X DPBS, DQ-BSA were chased for 2 hours in phenol red-free complete DMEM at 37°C. Following chase, cells were then loaded with 100µg/mL10kDa fluorescein (FITC) dextran for 30 minutes. FITC dextran was not chased and imaged immediately after washing and replacement with fresh pre-warmed phenol-red free complete DMEM. Live time lapse images were captured via spinning disk microscopy as described with LLOMe added on the stage after 2 minutes.

#### Magic Red-Cathepsin B

For use of Magic Red Cathepsin-B (MR-B) (ImmunoChemistry Technologies), MR-B stock was prepared as described in manufacturer’s protocol and diluted 1:1000 in complete DMEM. Cells were incubated with dilute MR-B for 20 minutes at 37°C. Cells were then sufficiently washed three times with 1X DPBS and replaced with fresh complete phenol red-free DMEM. MR-B loaded cells were equilibrated by incubation at 37°C for 5 minutes prior to spinning disk confocal microscopy. Change in cellular MR-B fluorescence over time was plotted relative to initial intensity.

### Western blotting

Samples containing 1.2X Sample Buffer (Genscript) were separated via SDS-PAGE using 4-12% Bis-Tris (Genscript) in 1X MES Running Buffer (Genscript) according to manufacturer’s instructions. Protein was transferred to nitrocellulose membranes (Bio-Rad) using the high molecular weight program of the Trans-Blot Turbo Transfer system (Bio-Rad) according to manufacturer’s instructions. Blocking was performed for 1 hour at room temperature with 100% LI-COR Odyssey Blocking Buffer. Primary antibody incubation (4°C, 16 hours) and secondary antibody incubations (22°C, 1 hour) were performed in 50% Odyssey Blocking Buffer/50% TBS + 0.1% Tween-20. Washes were conducted following primary and secondary incubations, 3 times 10 minutes, with excess TBS + 0.1% Tween-20. Blots were visualized and captured using the LI-COR Odyssey CL-X imaging system with LI-COR ImageStudio software. Densitometry analyses were performed using ImageStudio software.

### Rab7 activity assays

Preparation of bead conjugated GST-fusion proteins and GST-RILP based pulldown assays were performed as described in (Romero Rosales et al., 2009). GST fusion proteins (GST-stop and GST-RILP) were transformed and isolated from BL21 *E.coli* and conjugated to Pierce Superflow GST agarose (Thermo Scientific). To probe for Rab7-GTP, COS-7 cells expressing Em-Rab7^WT^ were treated with DMSO or 1mM LLOMe for 1 hour. Cells were collected by scraping in 1mL PBS containing Complete EDTA-free Protease Inhibitor (Roche) and pelleted. Cell pellets were resuspended in 200µL of GST Pulldown Buffer [20mM HEPES pH 7.4, 100mM NaCl, 5mM MgCl_2_, 1% TX-100, Roche Complete Protease Inhibitors] and incubated on ice for 15 minutes. Samples were then spun at 14,000rpm for 15 minutes at 4°C. Supernatant was collected in a clean tube and quantitated using Micro-BCA Protein Assay Kit. Excess GST-stop and GST-RILP bound beads (150µg per reaction) were pre-equilibrated in GST Pulldown Buffer. 50µg of input lysates were added to pre-equilibrated beads and rotated for 16 hours at 4°C. Beads were then washed four times with GST Pulldown Buffer and re-suspended in 35µL of 1.2X Sample Buffer before visualization by SDS-PAGE and Western blotting for GFP and GST. 5% inputs were loaded in parallel. GST-RILP pulldowns of Em-Rab7^WT^ were normalized to input levels of Em-Rab7^WT^.

### Em-Rab7 Fluorescence Recovery After Photobleaching (FRAP)

Stable Em-Rab7^WT^ or Em-Rab7^L8A^ NRK cells were imaged live at 37°C in complete phenol red-free DMEM using the Nikon Ti2-E inverted microscope with AX-R confocal microscopy system and Nikon elements software. To generate FRAP movies, images were collected for one minute prior to bleaching at 1 frame per 10 seconds using the AX-R resonant scanner. To bleach Em-Rab7+ endosomes/lysosomes, a 33.0625μm^2^ area was bleached for 10 seconds using the 488-laser line (laser power = 15) and the AX-R galvano scanner. Images were then collected post-bleach for 4 minutes, at 1 frame per 10 seconds, using the AX-R resonant scanner. All movies were processed identically following collection using Nikon Elements Denoising.ai and subsequent histogram matching bleach correction in FIJI.

### siRNA Knockdown of Rab7

NRK or stable EmRab7^L8A^ expressing NRK cells were lipofected using Lipofectamine 2000 with empty mCherry plasmid and siRNA targeting either a non-targeted sequence (5′-UGGUUUACAUGUCGACUAA, Dharmacon #D-001810-01-05) or rat Rab7 (5′-GACCAAGAACACACACGUA, Dharmacon #J-089334-10-0010). After 1 hour of exposure to complexes, medium containing complexes was exchanged for complete DMEM medium. Knockdown was allowed to progress for 4 days prior to either fixation or FRAP analyses. To quantify degree of knockdown, equivalent numbers of mCherry- and mCherry+ cells were selected from each field and measured for fluorescence intensity in FIJI. Raw intensity values were divided by the cell area to provide fluorescence/unit area values, which were then normalized to the mean of the mCherry-group.

### Starvation Experiments

For starvation experiments, COS-7 cells plated on coverslips were transfected with either pmRFP-LC3 or HA-V_1_G_1_. 24h post-transfection, cells were washed 1X with sterile 1X-DPBS then replaced with either complete DMEM medium or EBSS (Sigma #E2888). 20 hours post medium replacement, wells designated in the Torin-1 treatment arm were replaced with fresh complete medium containing 1µM Torin-1 (Sigma #475991), and treated for 4 hours. Following 4-hour treatment, all wells were fixed and the HA-V_1_G_1_ transfected cells were immunostained for endogenous Rab7. 24-hour EBSS and 4-hour Torin-1 treatments were selected due to their ability to produce expected responses in pmRFP-LC3.

### Image and Statistical Analyses

All statistical analyses were performed using either R Studio or Prism 9 software. All assumptions were deemed to be met before a selection of a statistical test. Normality of residuals was tested using the Shapiro-Wilk test. Specific tests used are denoted in figure legends and statistical significance was defined as a p value < 0.05. All t-tests conducted were unpaired and two-tailed unless otherwise stated. All image analysis was conducted with either Bitplane Imaris 9.3.1 or FIJI (NIH).

#### Quantification of Rab7 compartment volume and area

Due to a mix of puncta and apparent cytosolic distributions of Rab7, the surface generation tool in Bitplane Imaris 9.3.1 were utilized to define Rab7 compartments. Rab7 surfaces >0µm^2^ and <30µm^2^ or >0µm^3^ and <5µm^3^ were filtered for 2-D area calculations and 3-D volumetric analyses, respectively, which eliminated the cytosolic pool and heavily clustered compartments. We confirmed that area and volume bounds filtered the same population of compartments. The number of compartments per cell, average surface size, and average surface fluorescence intensity per unit area for the entire field were reported.

#### Quantification of Airyscan endosomal marker colocalization

Two marker colocalization was determined using the colocalization function in Bitplane Imaris 9.3.1. Images were equivalently thresholded before determining Pearson’s and Mander’s correlation coefficients.

#### Quantification of time-lapse changes in Em-Rab7 area

Em-Rab7 objects and DQ-BSA objects were defined across all timepoints using the region-growing object generation tool in Imaris 9.3.1. Equivalent thresholds were used for creation of DQ*hi* and DQ*lo* objects across movies and conditions, within each replicate, based on an average intensity of a sample of DQ objects. Em-Rab7 objects were then filtered into DQ*lo* and DQ*hi* compartments based on colocalization with created DQ-BSA objects. The average Em-Rab7 object area per field of view was then extracted for each time point and fitted using R-studio. N= 10 (DMSO) and 9 (LLOMe) fields of view from 3 independent experiments.

#### Quantification of fluorescent dextran de-quenching

De-quenching of total cellular Fluorescein (FITC) following LLOMe treatment (as in Fig. S1A-C) was determined using Imaris. Whole cell regions of interest were denoted and cropped. FITC dextran objects were then created using the region-growing object generation tool. The average FITC object fluorescence intensity at each time point for each cell was then extracted and normalized to the first frame. Cumulative averages of all movies for each condition were then plotted over time using Prism. De-quenching of Oregon Green dextran (OGdex) fluorescence (as in Fig. 5) was determined using FIJI. Images were first processed using a median filter with a radius of 2 pixels. A binary mask of Oregon Green compartments was created and added to the ROI manager partitioned as individual objects for single frames 2 minutes prior and 3 minutes following DMSO or LLOMe addition. Fluorescence intensities of Oregon Green and DQ-BSA were measured within each ROI at each time point. ROIs were then filtered into DQ*lo* and DQ*hi* compartments (threshold= 10,000 a.u., determined by random sampling of apparent DQ*lo/negative* compartments by eye) and Oregon Green fluorescence was compared within each DQ filtered group using a paired t-test between pre- and post-treatment time points.

#### Quantification of ALIX+ LEs and LYS

Total ALIX puncta and colocalization of ALIX puncta with Rab7 and CatB was determined using the region-growing object generation tool in Imaris 9.3.1. Individual objects were created for each ALIX, Rab7 and CatB compartment. MatLab dual and triple object colocalization Imaris plugins were utilized to determine colocalized objects with overlapping radius= 1µm. Colocalization is expressed as the percentage of total Rab7 only, CatB only, or Rab7/CatB dual positive objects that colocalize with an ALIX object.

#### Quantification of ALIX responsiveness in BAPTA-AM

ALIX puncta and intensity in the presence or absence of Ca^2+^ chelation was quantitated using FIJI. Images were first processed using a median filter with a radius of 2 pixels. A binary mask of ALIX space was created in FIJI and applied to the original image as ROIs. The number of ROIs and average measured intensity of the ROIs were then extracted and reported.

#### Quantification of HA-V_1_G_1_ compartments

HA-V_1_G_1_ puncta number, intensity, and area was calculated using FIJI. All images were first processed using a median filter with a radius of 2 pixels. To determine HA-V_1_G_1_ intensity on endogenous Rab7 space (Fig. 7B, Fig. S8E-F), a binary mask of Rab7 area was first created. The mask was refined by then eliminating the nucleus and highly clustered Rab7 perinuclear regions which are non-permissive to distinguishing individual compartments. The refined binary mask was applied to the original image as individual ROIs and the fluorescence intensity of HA-V_1_G_1_ was then measured. Measured HA-V_1_G_1_ intensity was then normalized to the Rab7 ROI area and reported. To determine the HA-V_1_G_1_ puncta number and area (as in Fig. 7E-F, Fig. S7D-E, Fig. S8C-D), a binary mask of HA-V_1_G_1_ was created and similarly refined by eliminating the nuclear and highly clustered perinuclear regions. The mask was then applied to the original image as individual ROIs and the total number and area of individual ROIs was determined. The number and average area of all HA-V_1_G_1_ ROIs was then reported. In starvation experiments specifically, we only selected cells with responsive V1G1 cells to analyze for Rab7 changes, as the starvation response was not homogenous (∼50% of transfected cells).

#### FRAP Quantification

FRAP movies were equivalently processed using Nikon Elements Denoising.ai followed by Histogram Matching bleach corrections in FIJI. Using FIJI, the fluorescence intensity over time within the bleached area and an equivalently sized ROI control area was measured. Measurements within the bleached area were affinely scaled by subtracting the minimum intensity within the bleached area intensity from the bleached and control area intensity. Average intensity values at each time point were then plotted. FRAP recovery curves, starting at t= 70s, were fitted using least squares fit, with a one-phase exponential association equation in Prism. t_1/2_ and plateau values of fitted lines only were reported. Y_max_ values were calculated as the intensity of the bleached area at t = 300s, and compared across conditions using an ordinary One-Way ANOVA.

#### Rab7 Tubulation Quantification

Em-Rab7 tubulation in MEFs was quantified manually in FIJI. Images across conditions were randomized and blinded. Tubules were manually traced using the line tool in FIJI using the Rab7 channel, and confirmed to contain dextran prior to addition to the ROI manager. Tubules were explicitly defined as lighter linear membrane extensions emanating from a darker source endosome. Homogenous streaks were not counted due to the possibility of these being highly motile compartments. Total tubule ROIs from each cell were then measured for their length. The number and average length per cell was then reported.

#### Quantification of CI-M6PR Puncta and Distribution

CI-M6PR puncta number and Rab7 intensity were quantified in FIJI. The CI-M6PR channel for all images was first processed using a median filter with a radius of 2 pixels, then masked. To determine the number of CI-M6PR compartments, the mask was refined to eliminate the nuclear and highly clustered perinuclear regions which are non-permissive to distinguishing individual compartments. The number of remaining individual CI-M6PR ROIs was then counted and reported. To determine the intensity of Rab7 on CI-M6PR space, the unrefined CI-M6PR mask was applied to the un-processed Rab7 channel and the fluorescence intensity of Rab7 was measured. The average Rab7 intensity across ROIs was then calculated for each cell, and subsequently divided by the total CI-M6PR area to provide a value of Average Rab7 intensity per unit CI-M6PR area.

To determine peripheralization of CI-M6PR in WT- or Q67L- COS-7 cells (Fig. 3N) or EEA1 (Fig. S2E-F), we utilized an ImageJ plugin to measure the distance to the nucleus, and intensity of CI-M6PR compartments. Distances were binned into the perinuclear area (closest 33%) and peripheral area (distal 66%). In brief, equivalently transfected cells were selected, the nucleus and outer edge of the cell was manually annotated based on phalloidin staining, and a Euclidian distance map was created from the nucleus to the outer edge. Then, using a duplicate of the CI-M6PR or EEA1 channel, compartments were defined via thresholding and added to the region of interest manager, before being added back to the source CI-M6PR or EEA1 channel and the Euclidian distance map to measure intensity and distance values for each object, respectively. Using R, objects were then binned according to distance and the total intensity of objects within each bin was determined.

## Data Availability

Data available upon request.

## Acknowledgements

This work was supported by NIH R01NS083378 and R21NS128756 awarded to B.W., NIH T32GM007267 awarded to the UVA MSTP (PI-Dean Kedes), NIH T32GM139787 awarded to the UVA Cell and Molecular Biology Training Grant (PI-Bryce Paschal), and NIH F30AG079497 awarded to R.J.M.

We acknowledge the Keck Center for Cellular Imaging for the usage of the Zeiss 980 imaging system for generation of Airyscan data (PI-Ammasi Periasamy NIH-OD025156), and the Casanova and DeSimone labs for the usage of the Nikon TE 2000 spinning disk and Nikon AXR confocal microscopes, in addition to their critiques and support. We also acknowledge the UVA Molecular Electron Microscopy Core (RRID:SCR_019031) for their assistance. We again thank Dr. Cecilia Bucci for generously sharing the HA-V_1_G_1_ plasmid. We thank Dr. David Castle, Dr. James Casanova, Dr. Christopher Bott, Jonathon Sewell, and Isaiah Swann for critical reading of the manuscript.

Model figures (Fig. 2A, Fig. 6A, Fig. S5A, Fig. 8K) were created with Biorender.com. Figure panels were created using InkScape software.

## Author Contributions

R.J.M., C.C.Y., and B.W. conceived and coordinated completion of the study. M.M. and S.R. conceived, collected data, and interpreted electron microscopy data. R.J.M. collected and performed data analysis on all data. R.J.M prepared figures and R.J.M and B.W. prepared the first draft of the manuscript. Manuscript editing was performed by R.J.M., M.M., C.C.Y, S.R. and B.W.

## Author Notes

Disclosures: The authors declare no conflicts of interest.

