## supplemental figures and legends for "Collapse of late endosomal pH elicits a rapid Rab7 response via V-ATPase and RILP"

**Supplementary Figure 1 LLOMe disrupts organelles leading to pH collapse, membrane repair, and lysophagy in NRK and MEF cells**

**(A)** Schematic of pH dextran experiment: lysosomes were dually loaded with 100µg/mL pH-sensitive 10kDa FITC-dextran and pH-insensitive 70kDa TMR-dextran. Lysosomal disruption with LLOMe de-quenches FITC fluorescence. **(B)** Representative time-lapse images of NRK cells loaded with dual dextrans and exposed to 1mM LLOMe at t=0 min (scale=10µm). **(C)** Change in FITC fluorescence of the experiment in (B) quantified over time and fitted linearly (n=9 movies per condition from 3 independent experiments, see Video 1)**. (D)** Primary mouse embryonic fibroblasts (MEFs) treated with vehicle (DMSO) or 1mM LLOMe for 5 minutes were immunostained for ESCRT-associated protein ALIX (scale=25µm). **(E)** Number of ALIX puncta per cell were quantified and compared using Welch's t-test (n= 24 FOV from 3 independent experiments, line = median, ****p<0.0001). **(F)** NRK cells were treated with vehicle (DMSO) or 1mM LLOMe for one hour and immunostained for galectin-3 (Gal3) (scale= 25µm). **(G)** The percentage of Gal3+ cells per field of view were quantified over time. A washout condition was included to show that cells will clear damaged lysosomes in NRK cells, similar to what is reported for other cell lines. LLOMe-treated time points were compared to vehicle (DMSO) using One-Way ANOVA (n= 30 FOV per condition from 3 independent experiments, n.s. = not significant, *p<0.05, **p<0.01, ***p<0.001, ****p<0.0001. Error bars indicate Mean+ SEM).

**Supplemental Figure 2 Degradative Rab7+ lysosomes leak luminal peptides in LLOMe**

Live imaging of $EmRab7^{WT}$-transfected NRK cells loaded with the cathepsin-B substrate MR-B for 20 minutes and treated with vehicle (DMSO) **(A)** or 1mM LLOMe **(B)** (scale= 5μm). The small dipeptide leaks out of degradative Em-Rab7+ lysosomes, which is quantified over time via non-linear one phase decay equation in **(C)** (n= 9 movies per condition from 3 independent experiments). **(D)** MR-B puncta number in $EmRab7^{WT}$-transfected NRK cells collected as still images prior to and following 5-minute LLOMe treatment (1mM), quantified in FIJI, and compared using Welch's t-test (*p<0.0001) (n=58 (DMSO) and 59 (LLOMe) cells captured live from 2 independent experiments, line = median). **(E)** Schematic of EEA1 objects binned into the perinuclear Area 1 (first 33% of cell area) and peripheral Area 2 (outer 66% of cell area), and measured for intensity of EEA1 in Area 1 **(F)** Percentage of total cellular EEA1 intensity in Area 1 was compared using unpaired two-tailed student's t-test (p =0.0779) (n=23 (DMSO) and 29 (LLOMe 1mM, 1h) cells from 9 Airyscan micrographs per condition, line = median)

**Supplemental Figure 3 Electron microscopy demonstrates a variety of altered endo-lysosomal compartments in LLOMe**
**(A)** Endosomal/lysosomal diameters in NRK cells treated with DMSO or 1mM LLOMe for 45min, measured manually on electron micrographs and compared using Mann-Whitney U test (n =275 (DMSO)/546(LLOMe) endosome/lysosomes from 2 independent experiments, ****$p<0.0001$, line = median). **(B)** Rare examples of enlarged compartments observed in DMSO conditions. **(C)** Numerous examples of frequently observed single membraned, lucent compartments in LLOMe, containing intraluminal vesicles (see also Fig.1G, yellow arrow). **(D)** Numerous examples of frequently observed single membraned, material laden compartments, containing apparent organelles, membranes and dense material (scale bars = 1 µm). **(E)** Examples of occasionally observed double membraned compartments, most likely to be autophagosomes (scale bars = 1 µm). **(F)** Examples of occasional irregularly shaped, single membraned, and lucent compartments (scale bars = 1 µm).

**Supplemental Figure 4 Early endosomes are preserved in LLOMe**

**(A)** Airyscan micrographs of NRK cells treated with vehicle (DMSO) or LLOMe (1mM, 1h), immunostained for Rab7 (green) and EEA1(early endosomes; magenta) and imaged by Airyscan microscopy (scale= 5µm, inset= 1µm). Yellow arrowheads point at EEA1+ Rab7+ transitioning EE/LEs which are not enlarged. Mander's correlation coefficients for **(B)** EEA1 intensity on Rab7 space (p = 0.7712) and **(C)** Rab7 intensity on EEA1 space (p= 0.5310) were calculated using Imaris and compared using unpaired two-tailed student's t-test (n= 9 FOV per condition from 3 independent experiments, Error bars = mean ± SD). **(D)** Individual live imaging traces of average cellular EmRab7$^{WT}$ object volumes over time (blue = DMSO 10 cells; pink = LLOMe 9 cells). These traces were combined for statistical analysis in Fig. 2 G and I.

**Supplemental Figure 5 $Ca^{2+}$ release from ruptured lysosomes is not responsible for alterations in Rab7 in LLOMe**

**(A)** Schematic for LE/LYS $Ca^{2+}$ manipulation including cytosolic $Ca^{2+}$ chelation with cell permeable BAPTA-AM and stimulated $Ca^{2+}$ release using the TRPML1 agonist ML-SA1.
**(B)** Representative micrographs of primary MEFs pre-treated (or not) with 10μM BAPTA-AM for 30 minutes and then treated for 5 min with DMSO or 1mM LLOMe. Cells were then fixed and immunostained for endogenous Rab7 (green) and ALIX (magenta) (scale = 5μm). Rab7 intensity and size still increased in +BAPTA-AM even though ALIX recruitment was abolished. Rab7 intensity levels were artificially matched for viewing. **(C)** Quantification of ALIX puncta intensity and **(D)** number per cell following LLOMe and BAPTA-AM treatments and compared using Kruskal-Wallis test (n = 40 cells per condition from 2 independent experiments, line = median, n.s. = not significant, *p<0.05, **p<0.01, ***p<0.001, ****p<0.0001). Similarly, to published results, chelation of cytosolic calcium with BAPTA-AM after LLOMe diminishes ALIX recruitment. **(E)** Representative stills from spinning disk microscopy (see Video 5) of Emerald-$Rab7^{WT}$ expressing NRK cells exposed to vehicle, 1mM LLOMe or 20μM ML-SA1 for 10 minutes (scale = 10μm). Activation of lysosomal calcium efflux by ML-SA1 does not lead to changes in Rab7+ endosomes. These data argue that lysosomal calcium does not drive Rab7 changes observed in LLOMe.

**Supplemental Figure 6 Membrane rupture is only observed in LLOMe but not $NH_4Cl$ or BAF-A1.**

**(A)** Widefield micrographs of primary MEFs treated with vehicle (DMSO, 1 hour), LLOMe (1mM, 1 hour), $NH_4Cl$ (20mM, 1 hour), and BafA1 (200nM, 2 hour) and immunostained for endogenous Rab7 (cyan), ALIX (yellow), and Gal3 (magenta) (scale = 10μm, inset= 2μm). ALIX recruitment occurs with minimal membrane permeabilization whereas Gal3 requires larger membrane disruption. Neither occurs with $NH_4Cl$ or BafA1. **(B)** Under same conditions as (A), MEFs were high-pressure frozen and processed for transmission electron microscopy. Representative transmission electron micrographs for each condition are shown. Scale bar = 5μm, n = 1 independent experiment. **(C-F)** Representative examples of single compartments from control and pH perturbation treatments in MEFs demonstrate similarities between frequently appearing endosome/lysosome compartments in LLOMe (D), $NH_4Cl$ (E), and BafA1 (F) treatments (scale = 1μm). Control (DMSO) is shown in C.

**Supplemental Figure 7 Rab7 is not essential for $V_1G_1$ recruitment to endosomes**

**(A-E)** Rab7 activation state does not impact HA-$V_1G_1$ recruitment. (A) Confocal micrographs of COS-7 cells transfected with GFP-$Rab7^{WT}$ and HA-$V_1G_1$ treated with vehicle (DMSO) or 1mM LLOMe for 30 minutes (scale= 10µm, inset= 5µm). (B) Confocal micrographs of COS-7 cells transfected with constitutively active (CA) GFP-$Rab7^{Q67L}$ and HA-$V_1G_1$ treated with vehicle (DMSO) or 1mM LLOMe for 30 minutes (scale= 10µm, inset= 5µm). (C) Confocal micrographs of COS-7 cells transfected with dominant negative (DN) GFP-$Rab7^{T22N}$ and HA-$V_1G_1$ treated with vehicle (DMSO) or 1mM LLOMe for 30 minutes (scale= 10µm, inset= 5µm). Quantification of the number (D) and average area (E) of peripheral $V_1G_1$ objects from (A-D) compared using Kruskal Wallis test (n= 51 (WT-DMSO), 62 (WT-LLOMe), 59 (Q67L-DMSO), 60 (Q67L-LLOMe), 52 (T22N-DMSO), 65 (T22N-LLOMe) equivalent dual transfected cells, n.s. = not significant, ***p<0.001, ****p<0.0001). **(F)** Rab7 is not required for HA-$V_1G_1$ recruitment. Widefield micrographs of untreated primary $Rab7^{fl/fl}$ MEFs transfected with HA-$V_1G_1$ and GFP or pCAG-GFP-Cre (yellow) for 3 days, and immunostained for endogenous Rab7 (cyan) and HA-tag (magenta). (scale = 20µm). **(G-I)** BafA1 causes endosome/lysosome pH collapse across cell types. Cells were loaded with 10kDa FITC dextran (green) and 70kDa TMR dextran (magenta) overnight, then chased for 4 hours prior to treatment with either vehicle (DMSO) or BafA1 (200nM, 2hr) followed by spinning disk confocal microscopy. All endosome/lysosomes were thresholded as ROI using the TMR channel in the entire fields of (G) COS-7 [DMSO-20, BafA1-22 FOV], (H) NRK [DMSO-20, BafA1-24 FOV], and (I) MEF [DMSO-22, BafA1-22 FOV] cells, and then analyzed for the ratio of FITC:TMR within ROIs. Compared using either unpaired, two tailed student's t-test (NRK) or Mann-Whitney U test (COS-7 and MEF) (****p<0.0001, from 1 independent experiment).

**Supplemental Figure 8 Cellular starvation leads to Rab7 accumulation on $V_1G_1$-positive compartments**

**(A)** Widefield micrographs of COS-7 cells transfected with pmRFP-LC3 and treated with either complete medium, EBSS (24hr), or Torin-1 (1μM, 4hr) demonstrating expected LC3 responses in this cell type at the times and concentrations of starvation conditions used (scale = 20 μm, inset = 5 μm). **(B)** Widefield micrographs of COS-7 cells transfected with HA-$V_1G_1$, treated with either complete medium, EBSS (24hr), or Torin-1 (1μM, 4hr), and immunostained for endogenous Rab7 and HA-tag. Only $V_1$-responsive cells are shown and analyzed (~50% total transfected cell population) (scale = 20μm, inset = 5μm, yellow arrowhead = representative $V_1G_1$ compartments). **(C)** Quantification of the number (*left*) and area (*right*) of $V_1$ puncta in (B). **(D)** Quantification of the mean intensity of endogenous Rab7 on $V_1$ space, per cell in (B). **(E)** Quantification of the number (*left*) and area (*right*) of Rab7 puncta in (B). **(F)** Quantification of the mean intensity of HA-V1G1 on Rab7 space, per cell in (B). (C-F) compared using Kruskal Wallis Test (n= 48 (Complete), 48 (EBSS), and 51 (Torin) equivalently transfected cells per condition, line = median, from 2 independent experiments). **(G)** Widefield micrographs of NRK cells lipofected with a mixture of mCherry plasmid and anti-Rab7 siRNA (magenta) for 4 days, and immunostained for endogenous Rab7 (green). Representative mCherry/siRab7+ cell (yellow arrowhead) and mCherry- (white arrowhead) are shown (scale = 20μm). **(H)** Endogenous Rab7 intensity per cell was quantified for mCherry- and mCherry+ NRK cells and compared using Mann-Whitney U test (n= 61 (mCherry-) and 54 (mCherry+) cells, ****p<0.0001, line = median, from 1 experiment). **(I)** Representative stills from confocal FRAP live imaging of mCherry+/EmRab7$^{L8A}$ NRK cells lipofected with mCherry and either non-targeting siRNA (siNT) or siRab7 under DMSO (10mins) or LLOMe (1mM, 10mins) conditions (n= siNT/DMSO (15 cells), siRab7/DMSO (15 cells), siNT/LLOMe (21 cells), siRab7/LLOMe (18 cells), from 2 independent experiments). Only EmRab7$^{L8A}$ fluorescence is shown for clarity. Bleached area indicated by white box (33.0625μm$^2$) with representative Rab7+ endosomes= yellow arrowhead (scale = 2μm). **(J)** FRAP recovery curves were plotted over time for siNT versus siRab7 under DMSO and LLOMe conditions. **(K)** Mean FRAP area intensity values were determined at t= 300sec ($Y_{max}$) and compared using One-Way ANOVA (n= as listed in (I), 2 independent experiments, line = median).

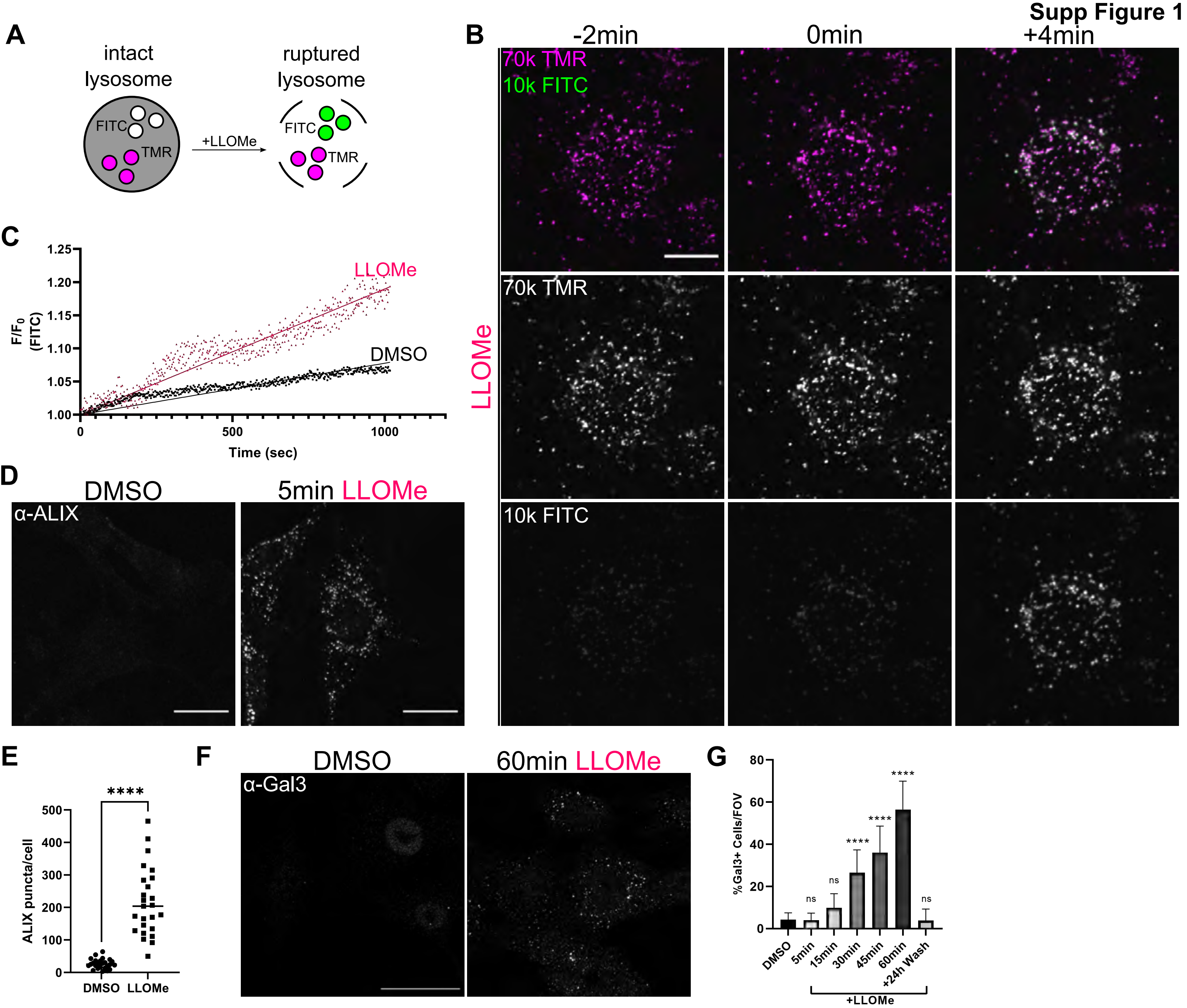

Supp Figure 1
A
intact lysosome
FITC
TMR
+LLOMe
ruptured lysosome
FITC
TMR
B
-2min
0min
+4min
70k TMR
10k FITC
LLOMe
70k TMR
10k FITC
C
1.25
1.20
1.15
1.10
1.05
1.00
F/F0 (FITC)
LLOMe
DMSO
0
500
1000
Time (sec)
D
DMSO
5min LLOMe
α-ALIX
E
****
500
400
300
200
100
0
ALIX puncta/cell
DMSO
LLOMe
F
DMSO
60min LLOMe
α-Gal3
G
80
60
40
20
0
%Gal3+ Cells/FOV
ns
ns
****
****
****
ns
DMSO
5min
15min
30min
45min
60min
+24h Wash
+LLOMe

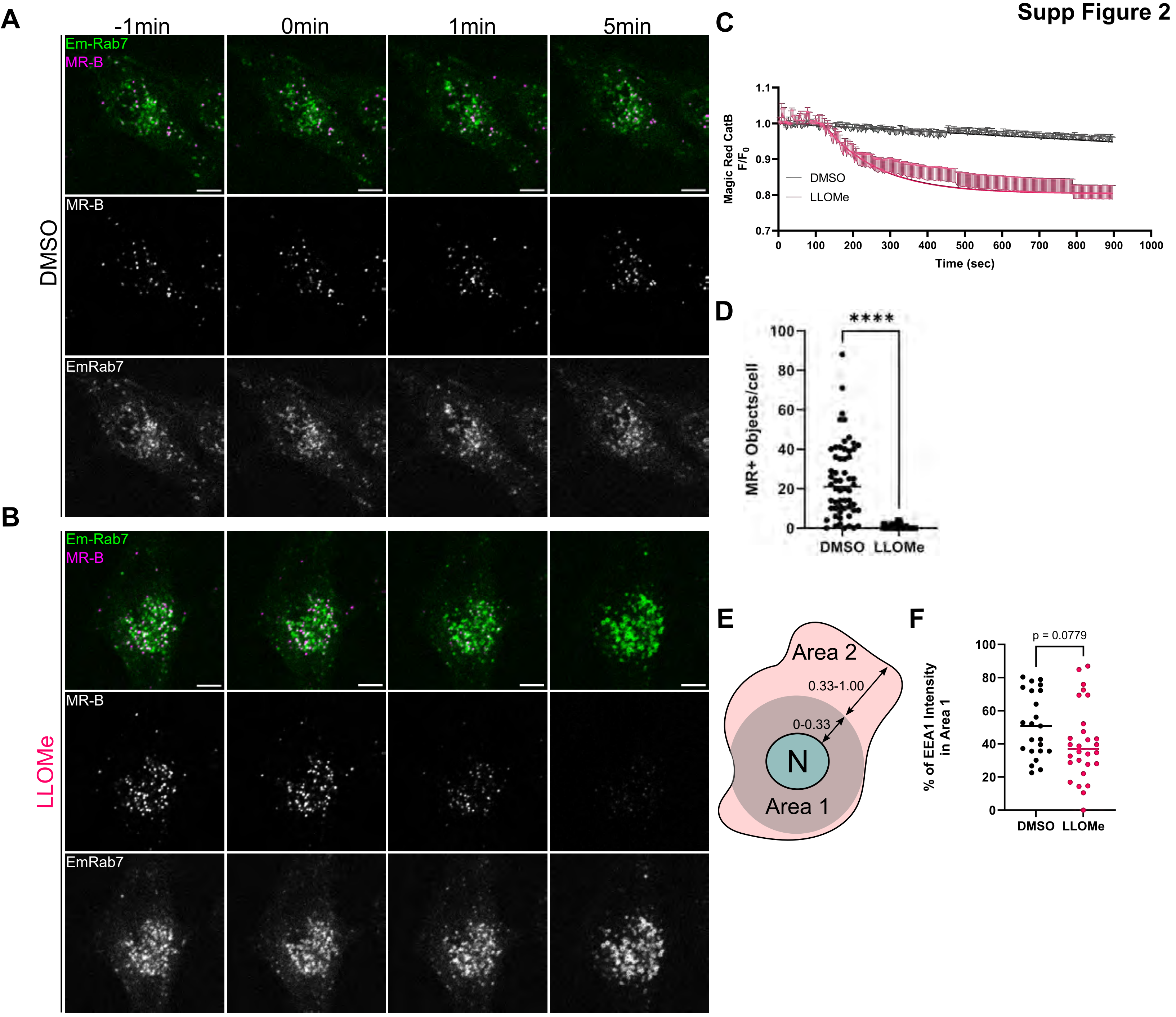
Supp Figure 2
A
-1min
0min
1min
5min
Em-Rab7
MR-B
DMSO
MR-B
EmRab7
B
Em-Rab7
MR-B
LLOMe
MR-B
EmRab7
C
Magic Red CatB F/F0
DMSO
LLOMe
Time (sec)
D
MR+ Objects/cell
****
DMSO
LLOMe
E
Area 2
0.33-1.00
0-0.33
N
Area 1
F
p = 0.0779
% of EEA1 Intensity in Area 1
DMSO
LLOMe

Supp Figure 3

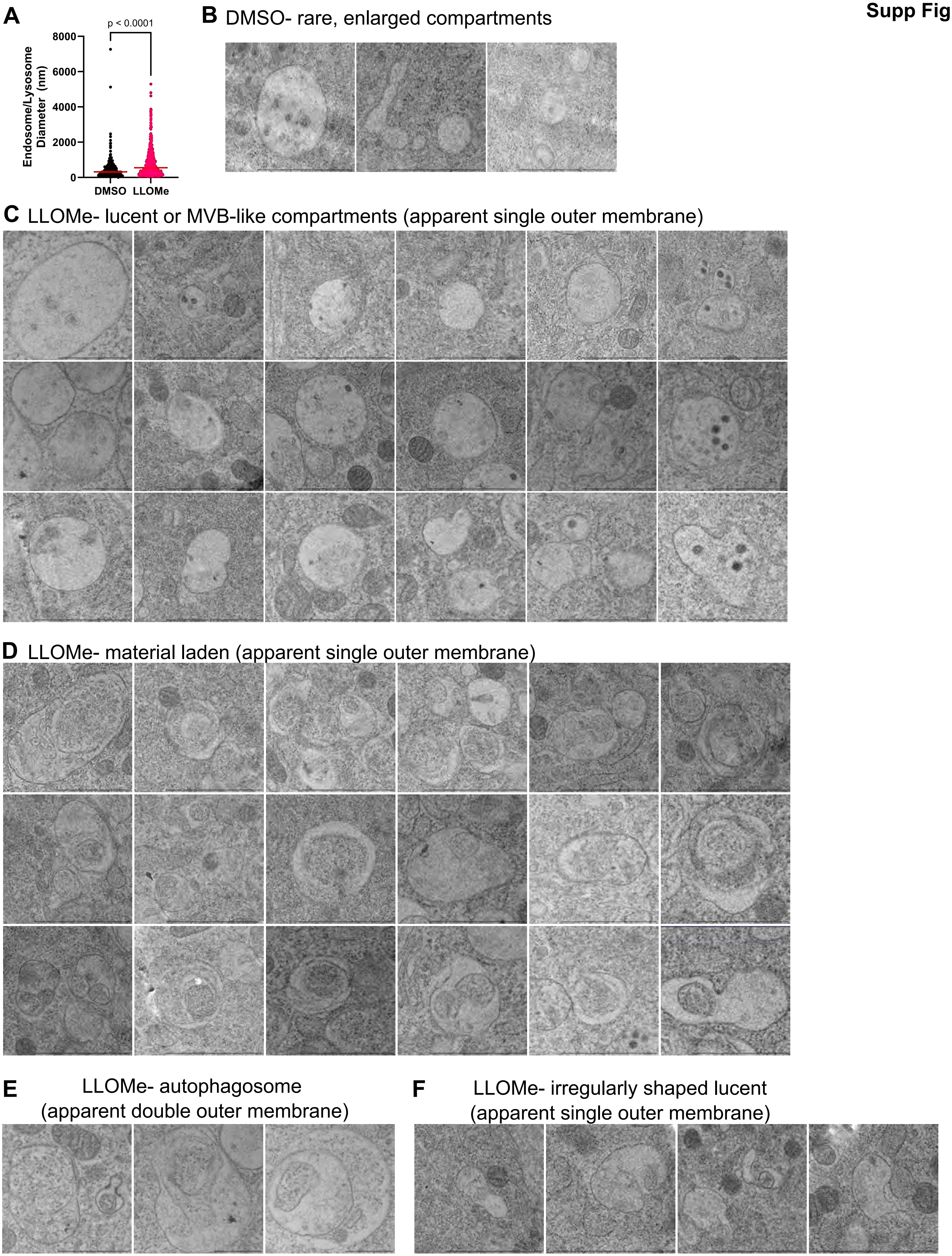

### Supp Figure 4

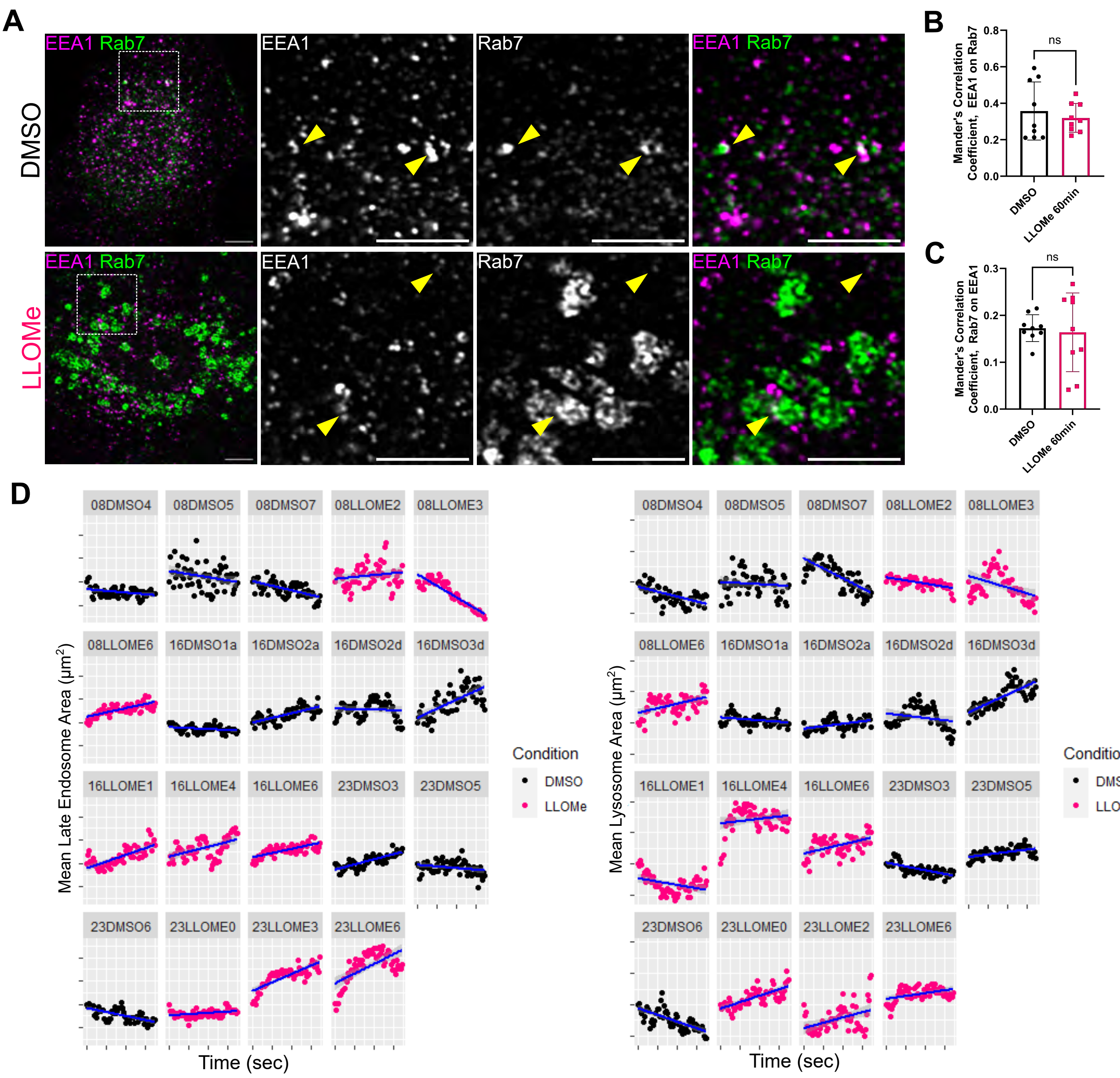

**Supp Figure 5**

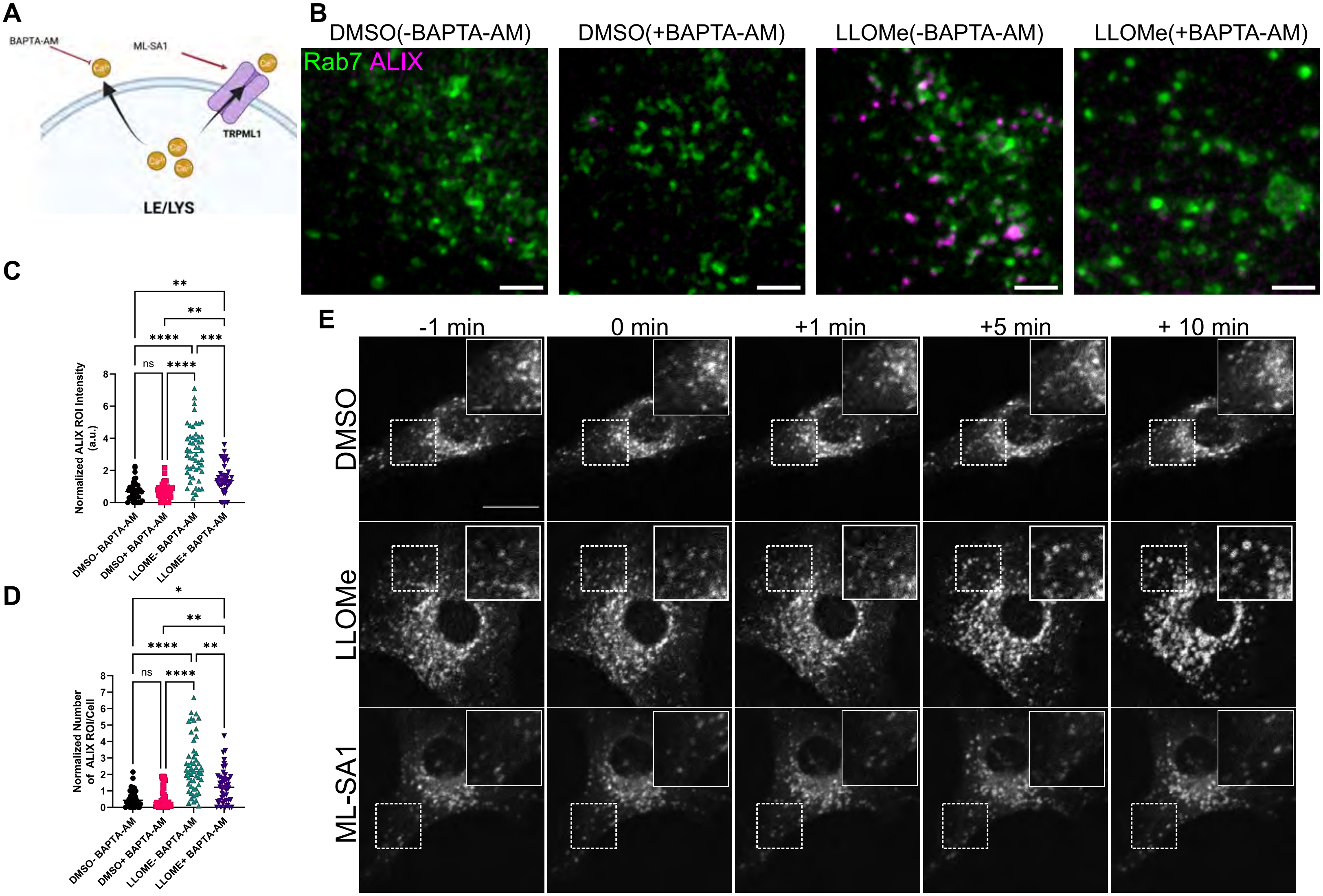

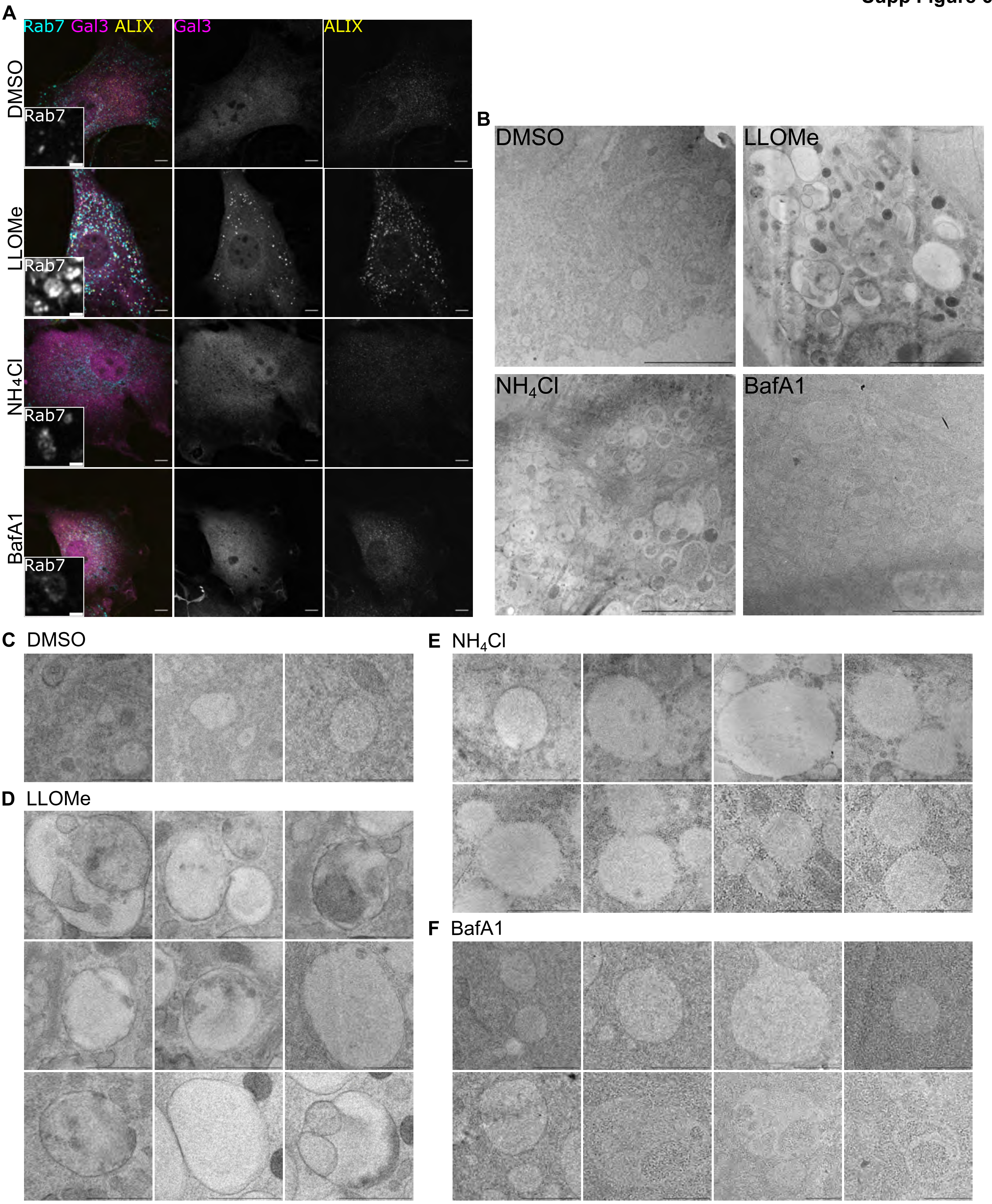
A
Rab7 Gal3 ALIX
Gal3
ALIX
DMSO
LLOMe
NH4Cl
BafA1
Rab7
B
DMSO
LLOMe
NH4Cl
BafA1
C DMSO
D LLOMe
E NH4Cl
F BafA1

Supp Figure 7

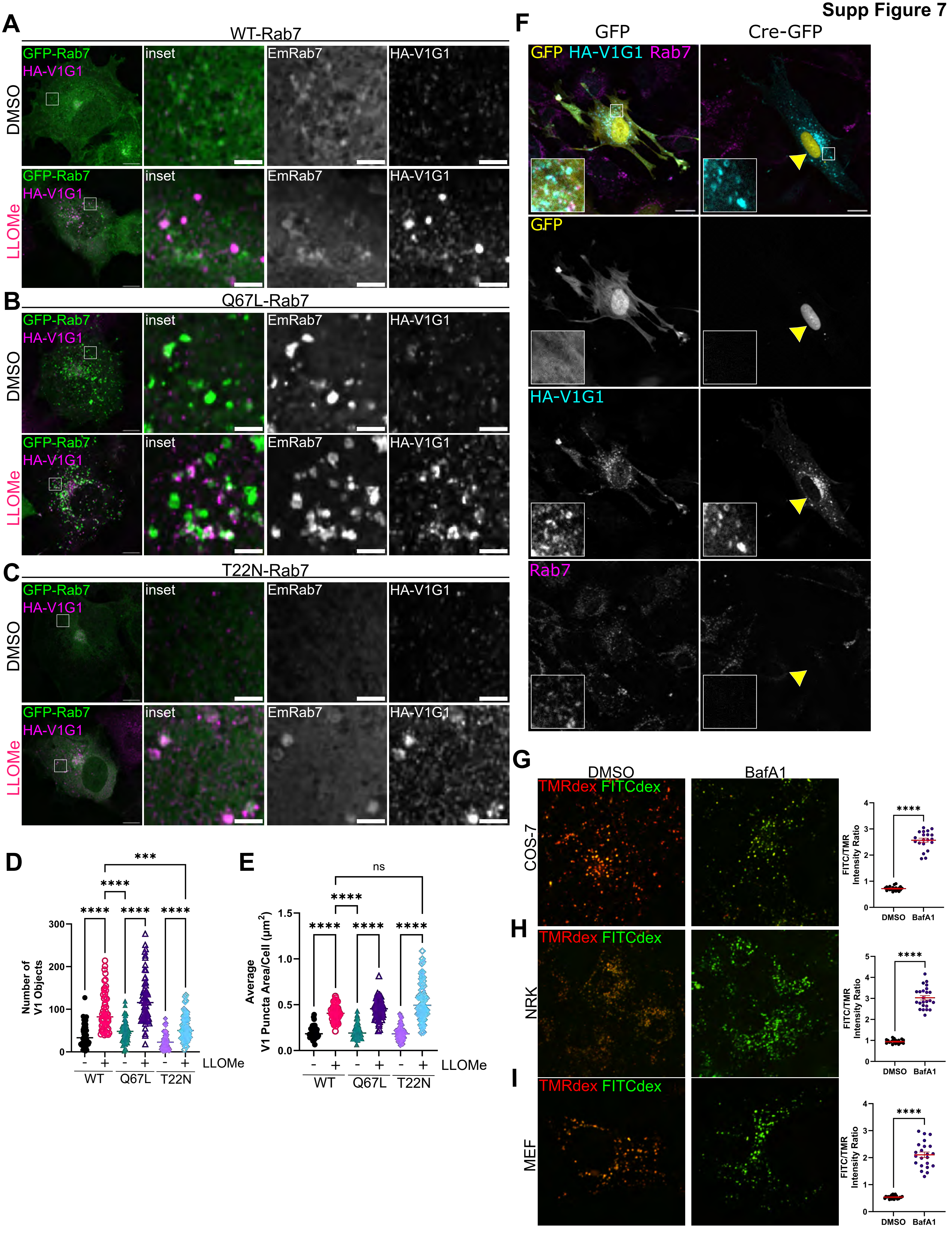

Supp Figure 8

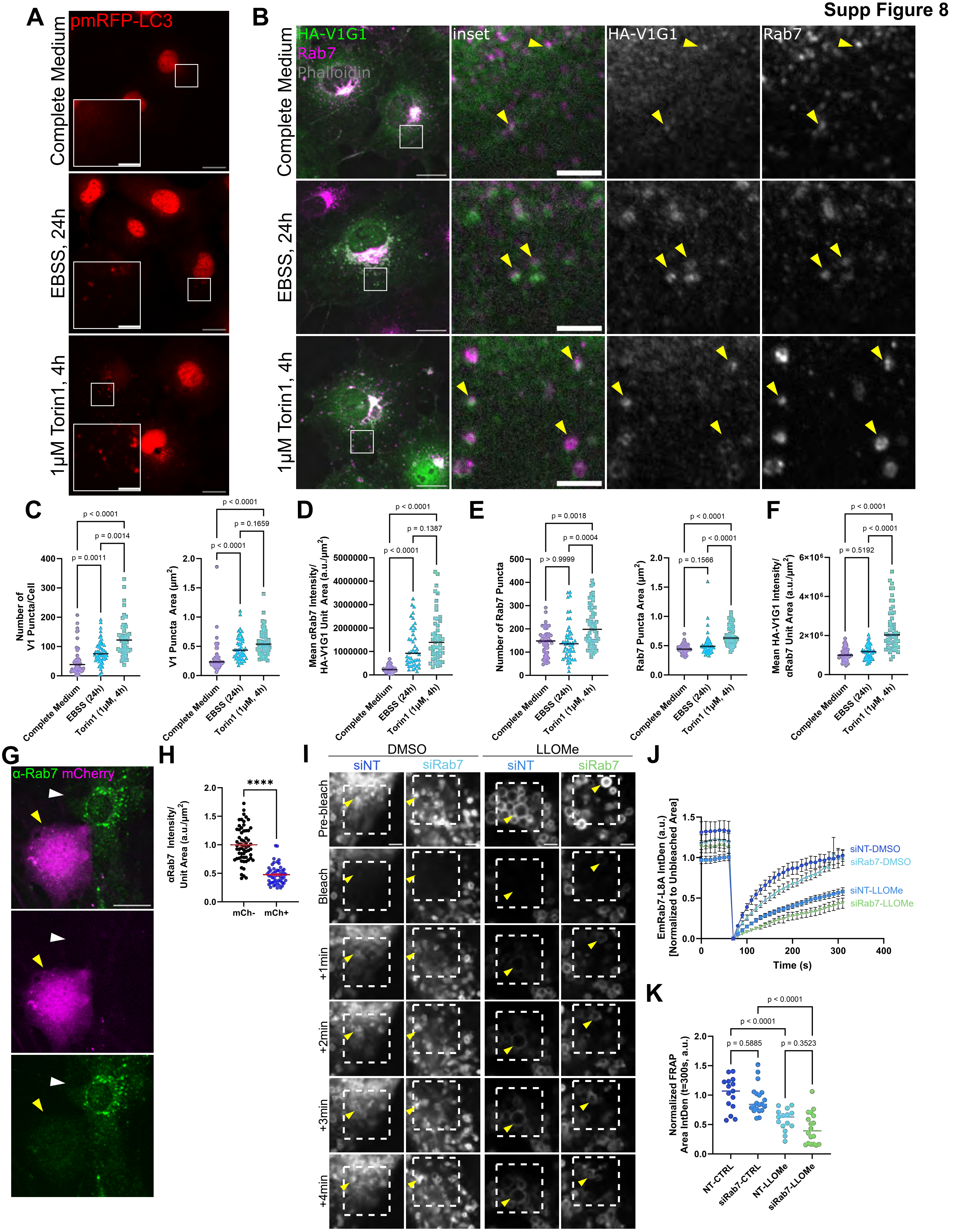
